# All-Optical Strategies to Minimize Photo-Bleaching in Reversibly Switchable Fluorescent Proteins

**DOI:** 10.1101/2025.01.20.633912

**Authors:** Guillem Marín-Aguilera, Francesca Pennacchietti, Andrea Volpato, Alessia Papalini, Abhilash Kulkarni, Niusha Bagheri, Guillaume Minet, Jerker Widengren, Ilaria Testa

## Abstract

Photo-bleaching is a general hurdle of fluorescence-based techniques that becomes even more severe in high-resolution microscopy that relies on prolonged, focused and complex illumination sequences. Strategies to reduce photo-bleaching require chemical modifications of the cell media, which often stave off physiological cellular conditions. Here, we outline an all-optical strategy to minimize photo-bleaching in reversibly switching fluorescent proteins (RSFPs), a class of probes used in several super-resolution and protein-multiplexing imaging techniques. By identifying the photobleaching pathways, we developed novel imaging schemes to increase the number of ON-OFF photo-switching cycles based on a designed modulation of the on-switching light or a co-irradiation with red-shifted light. By rationalizing the photo-cycle, we expand multiplexing strategies with RSFPs to high-spatiotemporal resolutions while maintaining the accuracy and recording longer time-lapse imaging of sub-cellular structures with both confocal microscopy and parallelized RESOLFT nanoscopy.

## Introduction

Reversibly switchable FPs (RSFPs) have advanced the field of fluorescence microscopy by enabling new super-resolution imaging^1–7^ and spectroscopic approaches^8,9^, as well as allowing for kinetic-based imaging multiplexing^10–13^. The switching mechanism in RSFPs involves a light-induced transition between two distinct molecular species, an emissive (on-state) and a non-emissive (off-state). The nature of the photo-switching is generally associated to cis/trans isomerization of the chromophore followed by a protonation/deprotonation^14–18^. Due to the stable off-switching state, it is possible to create enough sparsity to use them in single-molecule localization microscopy^5,19^ and super-resolution optical fluctuation imaging (SOFI)^20,21^, while the reversible nature of the switching mechanism makes them compatible with coordinate-targeted techniques, in particular, reversible saturable optical fluorescent transition^1^ (RESOLFT) and non-linear structured illumination microscopy^3^ (NL-SIM).

The diversity in shapes and speed of the off-switching kinetics associated with different variants has been used in imaging multiplexing at the cellular and subcellular levels^11–13,22^. In fluorescence-based spectroscopic techniques such as fluorescent anisotropy, the long lifetime of the RSFP’s ON-OFF states enables to extend the mass limit inherent to conventional fluorescence anisotropy and measure the rotational diffusivity of molecules and complexes with larger masses and hydrodynamic radius^8^.

Regardless of the approach, the ability of RSFPs to withstand several rounds of photo-switching cycles defines the resulting image quality. The progressive loss of the fluorescence upon cycling is called photo-switching fatigue and reflects the photo-bleaching profile of the fluorescent protein. In a broader sense, photo-bleaching results from the interplay between the local nano-environment of the probe and its complex photo-physics upon excitation, thus, efforts to mitigate photo-bleaching in microscopy strived to optimize the conditions on those two fronts. Regarding the former, multiple studies have shown that refining the composition of the imaging solvent improved the probe’s photo-stability^23–25^. Fusing enhancer nanobody to proteins of the rsGreen family increased their brightness and photostability ^26^. At the same time, the development of RSFPs as fluorescent markers has been guided by recursively introducing mutations that favour faster and more robust photo-switchers.

Studies in EGFP identify the triplet state as the gateway for photo-bleaching, being the leading pathway for oxidative chemistry in the chromophore^27^. Recent studies have characterised the triplet lifetime of rsEGFP2 as well as its absorption spectrum which presents a peak around 488 nm^28^, the same wavelength used to elicit the fluorescent photons. The properties of the triplet state can be leveraged to reduce the photo-bleaching of FPs^29^. Given that the long-lived triplet state of FPs has a red-shifted spectrum compared to the emissive form, co- illuminating in this spectral window can promote the depopulation of the dark state back to the ground-state singlet. The major photo-bleaching pathway becomes less efficient enabling longer time-lapse imaging^29^. Although optically-induced repopulation of the emissive state from the triplet has been known for years^30,31^, and even used in imaging with organic dyes^32,33^, it had not been applied to photo-bleaching reduction until recently^29^. Nonetheless, the range of fluorescence excitation intensities where the red-induced photo-bleaching reduction is predicted to be significant for the reported FPs is far from the power densities of scanning techniques, especially the one typically used in super-resolution microscopy.

Here, we characterize the main pathways behind photo-switching fatigue in RSFPs, which foster new prediction and implementation strategies to minimize photo-bleaching. This comprehensive understanding of the photo-cycle allows us to maximize the fluorescence output of rsEGFP2 by modulating the on-switching dose delivery.

Moreover, we develop a photo-switching model for rsEGFP2 that accounts for different sources of photo-bleaching and points to the optically induced depopulation of a dark state as a viable mechanism for photo-switching fatigue reduction at illumination intensities compatible with confocal and super-resolution imaging. The dark state is populated from the emissive species upon fluorescence excitation and can be recovered upon exposure to lower-energy red light. By co-illuminating the probes with red-shifted light to fluorescence excitation, we show a decrease in photo-switching fatigue for a library of green-emitting RSFPs. Based on this photophysical investigation, we propose a strategy to prolong confocal and super-resolution time-lapse imaging. The new imaging scheme features patterned light at different wavelengths and enables extended time-lapse imaging of light-sensitive organelles.

## Results

### Characterization of photo-switching fatigue in rsEGFP2

To study the fatigue in RSFPs we focused on the negative photo-switcher rsEGFP2^34^. This FP is a common marker for super-resolution imaging^34–36^, multiplexed bio-imaging^11,12^ as well as novel fluorescence anisotropy methods^8^. The photo-cycle of rsEGFP2 has been extensively studied by means of spectroscopy and crystallography methods^34,37^ and the intermediate steps of the photo-cycle and their rates are well-reported in the literature^8,14–16^. Using this information, we built a quantitative model for rsEGFP2 including photo-bleaching trends under a regime of illumination relevant for conventional and super-resolution imaging. Importantly for rsEGFP2, and in general for all RSFPs, the off-switching provides a competitive channel to the triplet formation via inter-system crossing (ISC), which combined with the spectral shift between on and off state inherently shields the fluorophore from the adverse effects of prolonged excitation and decreases the probability of triplet-state accumulation. This is crucial when comparing the photo-switching fatigue, and therefore bleaching, in RSFPs to non-switching probes. Given a constant illumination the latter will form a steady-state triplet population during excitation, while in RSFPs, the triplet population will relax over time as a result of the off-switching process limiting the photo-bleaching to a specific time interval.

We homogenously illuminated a region (20×20 μm^2^) of a thin layer of rsEGFP2 embedded in polyacrylamide (PAA) gel with consecutive on-and-off photo-switching rounds. After each excitation cycle, the emitted fluorescent signal is captured by the camera and the photo-switching fatigue curve is built (Figure 1a). The photo-switching fatigue curve for rsEGFP2 is characterized by two components, i) an initial drop, a sharp loss of the fluorescent signal in the first few cycles, and ii) a steady decrease in the signal upon thousands of cycles, which we refer to as fatigue fraction.

**Figure 1.**
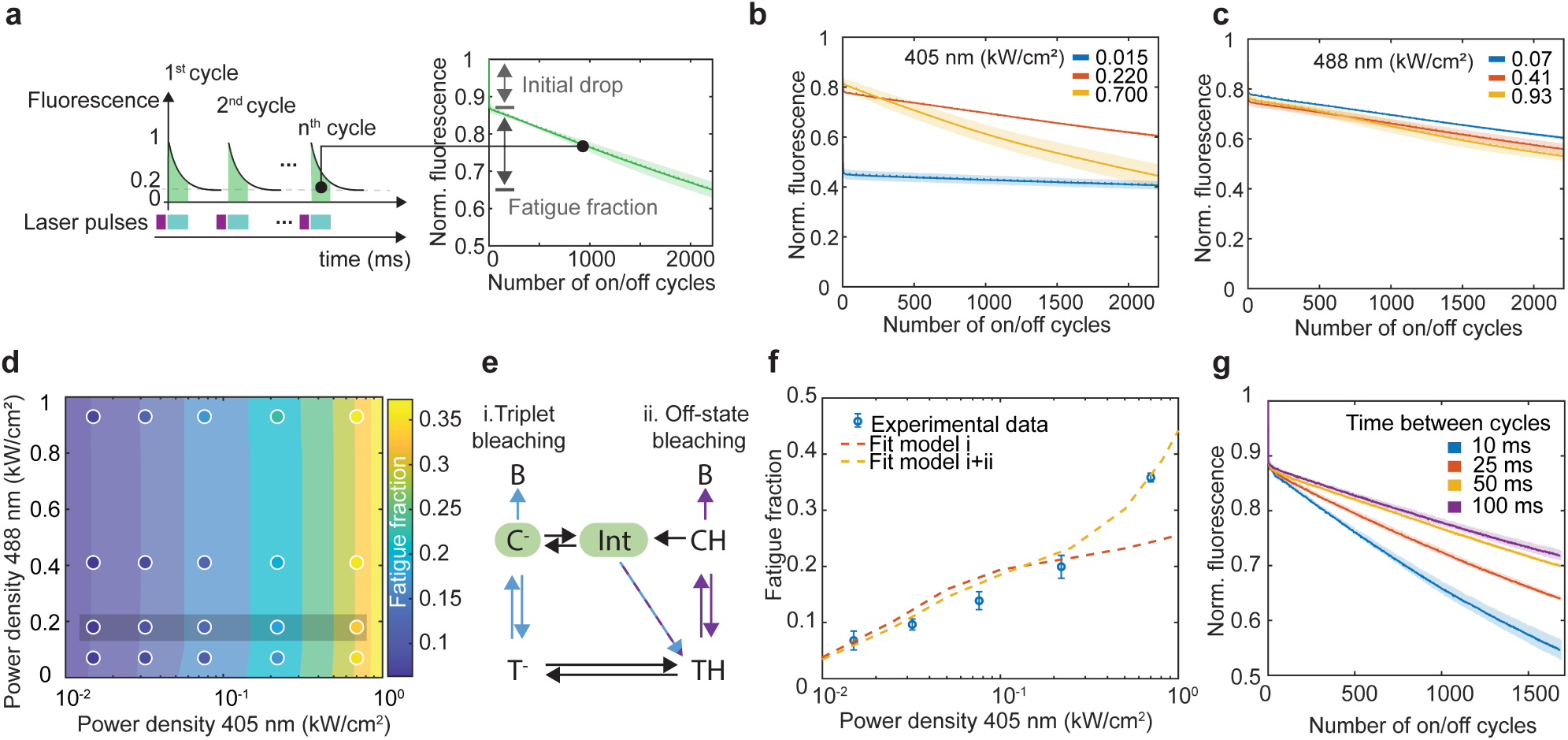
Photo-switching fatigue characterization and modelling. (a) Schematic of a photo-switching fatigue experiment. rsEGFP2-embedded PAA gel is repeatedly illuminated with 1ms UV-Vis (405 nm) and blue light (488 nm) for the time needed off- switch the fluorescence to 20%. The integrated fluorescence signal in each cycle is plotted against the number of cycles to generate the fatigue curve (mean ± σ of at least 3 measurements). (b, c) Photo-switching fatigue at different 405 nm (b) and 488 nm (c) illumination doses. (d) Photo-switching fatigue experimental data for a set of 405 and 488 nm power densities (dots) overlayed on top of the computed fatigue fraction from simulations. The colour bar indicates the magnitude of fluorescence loss between the 4^th^ and 2000^th^ cycle for every set of experimental conditions. (e) Kinetic scheme for rsEGFP2’s photo-cycle. The states are labelled distinctively depending on their conformation, Cis or Trans conformation, and their protonation state, H or ^-^. Black arrows indicate thermodynamically driven transitions while colored arrows indicate light-induced transitions. The colours of the arrows represent the most efficient wavelength to drive a given transition. (f) Comparison of bleaching models to the experimental data. In orange, only bleaching deriving from triplet state formation upon fluorescence excitation is considered. In yellow, the photo-switching fatigue is modelled considering two pathways, one stemming from the triplet and the other from 405 nm associated photo-damage. The experimental data corresponds to the data points in the shaded grey area in (d). (g) Effect of thermal relaxation on photo-switching fatigue by monitoring the photo-switching fatigue at increasing dark times between consecutive cycles.

We further studied the loss of fluorescence as a function of the illumination dose at 405 and 488 nm (Figure 1b-c). An increase in the 405 nm light dose decreased the observed initial drop, corresponding to a higher fraction of on- switched fluorophores per cycle. Similarly, the fatigue fraction also increased with higher 405 nm illumination doses, while displaying a minor dependency to 488 nm. Thus, we focused on investigating the fatigue fraction across nearly two orders of magnitude in power density for both illuminations (15 – 700 Wcm^-2^ for 405 nm and 70 – 930 Wcm^-2^ for 488 nm). As shown in Figure 1d, the data reports the 405 nm light to be the main driving force for increased photo-switching fatigue in rsEGFP2.

We rationalized the experimental data recorded at different illumination conditions for rsEGFP2 through a kinetic model including effective bleaching pathways (see Supplementary Note 1). We considered the photo-cycle of rsEGFP2 as a 5-state model (Figure 1e) where the photo-switching reaction occurs from a light-induced isomerization followed by a protonation/de-protonation process. The thermodynamically stable *cis anionic* (*C^-^*) form is also the emissive form of the chromophore. From there, the on-to-off photo-switching occurs via an excited state isomerization (cis-to-trans) reaction, followed by a ground state process to the *trans neutral* (*TH*) form^38,39^, with a characteristic time of ∼ 50 μs at physiological pH as revealed by high-power time-resolved off-switching experiments**^8^**. To recover the *C^-^* form of the chromophore, the off-to-on transition is efficiently triggered by UV light – typically 405 nm – from *TH*. Similarly, the excited state isomerization (trans-to-cis, in this case), to the *cis neutral* (*CH*) form, is followed by the deprotonation of the chromophore. As previously reported^14–16^, the off-to-on photo-switching process can be described as a cascade of intermediates at different time scales, of which an on-like protonated intermediate, *Int*, with a lifetime of ∼1 ms is relevant for the time scale under investigation (see Supplementary Note 2,3).

We incorporated the triplet-mediated photo-bleaching pathway in our modelling of rsEGFP2 (*i* in Figure 1e) and simulated the fatigue fraction response for a set of experimental conditions (405 nm power density ranging from 15 – 700 Wcm^-2^ and 488 power density at 180 Wcm^-2^). As shown in Figure 1f, the experimental data are not well supported by the simulation for high 405 nm power densities (PD > 300 W/cm^2^). The experimental behaviour, however, could be better described with an additional bleaching pathway stemming from the off-state (*ii* in Figure 1e), which will be mainly driven by 405 nm illumination (see Supplementary Note 4). This model can reproduce the increase in fatigue at high 405 nm illumination (Figure 1f) matching well the experimental data. The need for an additional bleaching channel is further supported by the dependence of the fatigue on the time between sequential cycles, giving the triplet population progressively more time to relax (Figure 1g). Increasing the dwell time from 10 to 100 ms resulted in ∼20 % photo-switching fatigue recovery after 1600 cycles (Figure 1g). However, the time dependence of such relaxation exceeds the expected dark state lifetime, both reported^28^ and measured in our data. This observation suggests that the triplet state acts as a precursor for slower photo-chemical processes that lead to photo-bleaching. Moreover, we observed that increasing the dark time by an order of magnitude does not result in the absence of photo-switching fatigue indicating that not all the irreversible fluorescence loss stems from a single mechanism.

### Maximizing ON-OFF cycles in rsEGFP2

The global photon budget of each RSFPs is set by the combination of the initial drop of the fluorescence and the number of cycles it can undergo. Based on the outlined model, we investigated strategies to maximize the detected fluorescence photons by modulating the illumination.

The initial drop appears as a sharp loss of fluorescence in the first few cycles of a photo-switching fatigue curve and is a consequence of the kinetics of the on-switching mechanism in rsEGFP2^40^. Its amplitude is defined by the deprotonation time and the absorption properties of the intermediate state (see Supplementary Note 3). At pH = 7.5, rsEGFP2 is expected to have 20% of the fluorophore’s concentration trapped in the *TH* (see Supplementary Note 2,3). The building up of the *TH* state can be altered by reducing the probability of the *Int*-to-*TH* transition. This can be achieved by preventing the accumulation of the intermediate state. In that regard, we investigated how the modulation of the on-switching dose, given a fixed total energy, influences the fluorescence output. The same 90 mJ/cm^2^ was delivered to the sample progressively increasing the illumination time (0.3 – 20 ms) while decreasing the 405 nm power density (300 – 4.5 W/cm^2^) as shown in Figure 2a. The fluorescence is enhanced up to 93% of the total expected signal (1^st^ illumination cycle with no 405 nm applied) when the activation dose is stretched over time to 20 ms and a corresponding 405 nm power density of 4.5 W/cm^2^ as shown in Figure 2a. Our simulations confirm that this is a result of preventing the quasi-equilibrium among *TH*, *CH* and *Int* (see Supplementary Note 5) and, therefore, a more efficient turnover between the on and off-states upon 405 nm irradiation. Altogether, we found that the initial drop observed in the photo-switching fatigue curves is not irreversible and can be efficiently recovered by modulating the way to deliver the on-switching light.

**Figure 2.**
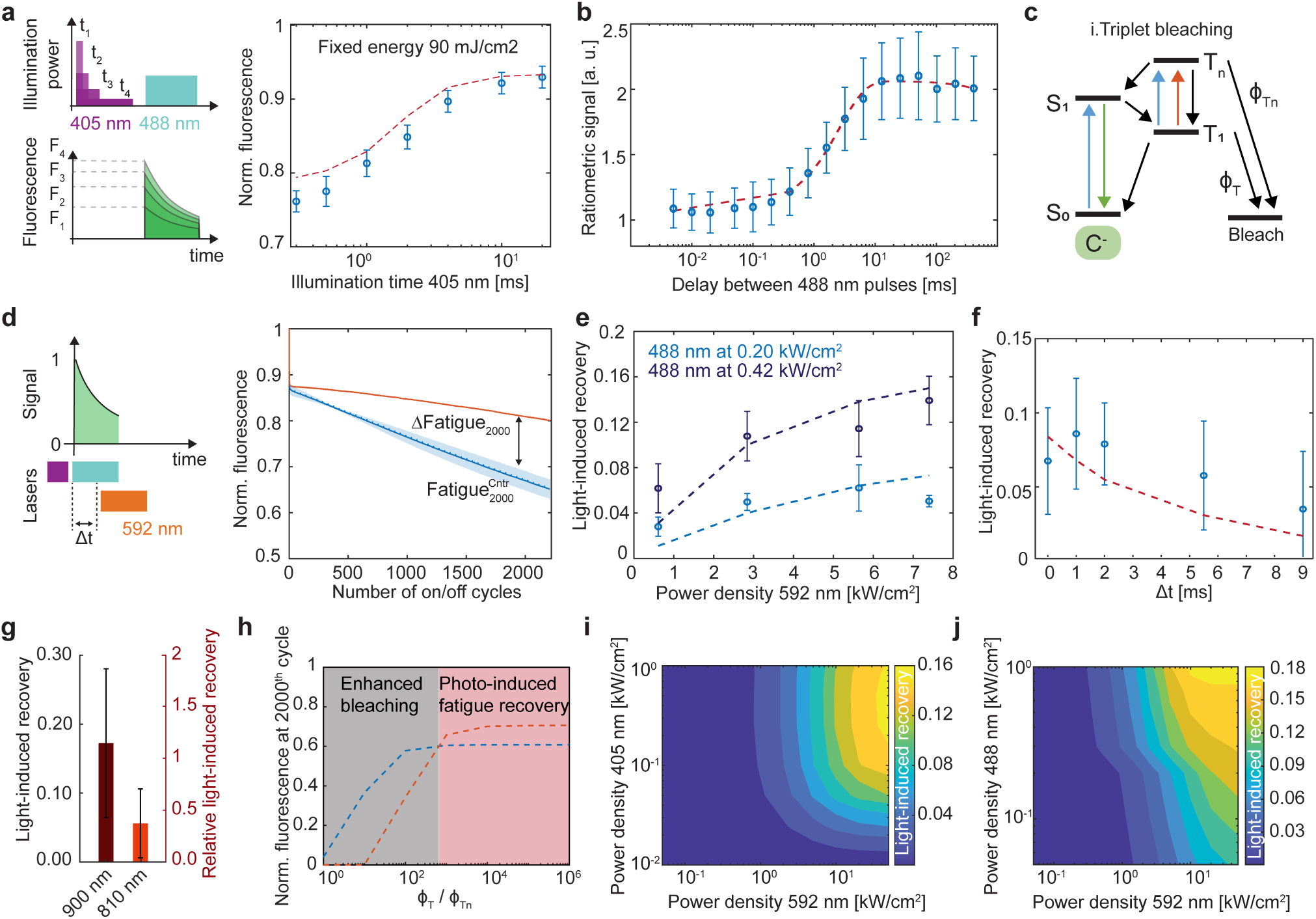
Optical strategies to minimize photo-bleaching in RSFPs. (a) Schematic of the on-switching modulation experiment, where the energy dose was kept constant at 90 mJ/cm^2^ and the 405 nm power density and illumination times progressively changed. The experiment was repeated over 4 different areas of a rsEGFP2- embedded PAA gel (mean ± σ). The input parameters for the simulated curve were the same as the experimental parameters. (b) Ratiometric signal as a function of the delay between consecutive 488 nm pulses to estimate the relaxation time of the dark state back to the emissive state (τ_1_ = 2.59 ± 0.20 ms). (c) Schematic of the proposed triplet bleaching pathway. The triplet absorption can be promoted upon blue and red illumination. Bleaching is considered from both the triplet ground (T_1_) and triplet excited state (T_n_). (d) Photo-switching fatigue curves with and without 592 nm co-illumination. To parametrize the recovery effect of the additional wavelength, we compared the fluorescence signal after 2000 frames with 592 nm to a control curve for each experiment. (e, f) Fatigue recovery dependence at increasing 592 nm power density (e) at two different 488 nm light doses and delay between 488 and 592 nm pulses (f). (g) Light-induced recovery dependence to different wavelengths in NIR spectral region. The power densities used for every red-shifted wavelength were ∼ 39 kW/cm^2^ at 900 nm and 48 kW/cm^2^ at 810 nm. (h) Fluorescence signal after 2000 cycles for different balances of the triplet bleaching channels. The recovery effect from the 592 nm co-illumination will be modulated by the relative strength of the triplet bleaching channels. Our recovery data suggest rsEGFP2 resides in the green-shaded area (i) Simulated light-induced recovery for a range of 405 and 592 nm power densities. For this simulation, the 488 nm dose was kept constant at 1.2 ms illumination time and 200 W/cm^2^. (j) Simulation of fatigue recovery as a function of the 488 and 592 nm power densities. An increase of the 592 nm dose shows an enhanced recovery for all 488 nm power densities. Important to note that the simulations were carried out at a constant 488 energy dose – power density * illumination time –.

To prevent photo-bleaching, reverse inter-system crossing (RISC), i.e. the backward process from the triplet state to the lowest singlet excited state, was shown to be optically modulable and helped to increase the brightness of organic dyes^30^. Since the protein scaffold shields the chromophore from the immediate local environment, the triplet lifetime in FPs is relatively long (∼ 5 ms for EGFP^27^) and RISC transition can be triggered with lower light doses compared to dyes (0.1-40 MWcm^-2^ for Eosin Y^30^, respect to ∼kWcm^-2^ for GFP-like proteins^29,41^). We measured the time evolution of a dark state relaxing in the μs-ms window by monitoring the increase in the integrated fluorescence signal between two 488 nm pulses separated by a variable delay. We observed a build-up on the signal of the second pulse with a raising time of ∼ 2.5 ms (Figure 2b and Supplementary Note 6). Since the thermal recovery rate for rsEGFP2 (thermal relaxation from *TH* to *C^-^*) has a timescale of several hours^37^, the increase in the concentration of the fluorescent emissive state is likely to originate from the relaxation of a dark state created upon excitation, i.e. the triplet state. The timescale of such relaxation is comparable to the reported triplet state lifetime in rsEGFP2 at 260K^28^.

To see if and to which extent the optically-induced RISC could help mitigate the photo-switching fatigue of RSFPs, we used the same experimental framework of the previous section with a modified pulse scheme which included red co-illumination (Figure 2d). We used 592 nm light, where the absorption cross-section for the triplet state of rsEGFP2 was reported to be ∼20 % of the peak at 900 nm at 100K^28^. The effect of the red-shifted wavelength was studied as a function of the power density (∼0.5-8 kWcm^-2^) as well as the delay time between the 488 and 592 nm pulses. The effect was quantified by comparing the difference in the fluorescence signal after 2000 cycles for a set of experimental conditions to a control fatigue curve, which is recorded without 592 nm illumination (Figure 2d). To provide further insights into the photo-switching fatigue mechanism, we fitted our experimental data to the newly developed model (see Supplementary Note 1) which covers the switching mechanism of rsEGFP2, the bleaching pathways described in the previous section as well as the spectroscopic properties of the triplet state, parametrized from the reported data in this manuscript and published studies^28^.

As we introduced the 592 nm co-illumination in the pulse scheme we observed a reduction in the photo-switching fatigue (Figure 2e), which increased at higher power densities up to 15%, suggesting that an absorption process is behind the recovery. We also observed a correlation between the magnitude of the recovery and the fluorescence excitation dose, with the light-induced recovery being more prevalent at higher 488 nm power densities (Figure 2e). Our data shows greater photo-switching fatigue at higher 488 nm power densities without 592 nm co- illumination while the red-shifted co-illumination restored the fluorescence signal to similar values for both 488 nm power densities (see Supplementary Note 6) suggesting the maximum recovery was reached. The magnitude of photo-switching fatigue recovery will be determined by: i) how big is the fatigue upon cycling at a given set of 405 and 488 nm power densities and ii) how much of the bleaching can be recovered by the red-shifted dose. In the same line, we characterized the photo-switching fatigue recovery as a function of the delay between 488 nm and 592 nm pulses (Figure 2f). Our results indicate that the magnitude of the recovery decays in the milliseconds time range which is compatible with the lifetime of the dark state characterized in Figure 2b.

Moving the red illumination closer to the peak of the triplet spectrum at 900 nm we expect an increase in the recovery. We experimentally validate the prediction by co-illuminating the RSFPs with Near Infra-Red light (NIR). The NIR co-illumination was tested for two wavelengths: 810 nm where the triplet state absorption coefficient is close to that at 592 nm^28^ and 900 nm, which is the peak of the triplet absorption spectra^28^. The recovery at the peak is ∼ 3 times higher than at the shoulder represented by the 810 nm. This is due to a higher triplet state absorption and lower cross-talk with other bleaching pathways, such as the aforementioned off-state bleaching (Figure 2g and Supplementary Note 7).

Our findings indicate that the photo-switching fatigue recovery stems from an optically-induced depopulation of the triplet state. Upon fluorescence excitation, both the triplet ground and excited states contribute to photo- bleaching (Figure 2c). Co-illumination with a red-shifted wavelength (592 nm) reduces the triplet state population, lowering the probability of photo-bleaching. The proposed kinetic model incorporates effective bleaching quantum yields for ground and excited triplet states – Φ_T_ and Φ_Tn_ respectively – (Figure 2c). The addition of a bleaching path from *T_n_* was necessary for reproducing the experimental fatigue curves, to not underestimate the experimental photo-bleaching. Such a pathway is seemingly relevant since the triplet state of rsEGFP2 presents an absorption peak at 488 nm^28^, and the triplet state absorption has been identified as a photo-bleaching promoter in EGFP^29^.

Using simulations, we studied how the balance between photo-bleaching pathways via the triplet state impacts photo-switching fatigue. In the absence of 592 nm co-illumination (Figure 2h, blue line), if both pathways have similar quantum yields, the fatigue fraction increases sharply compared to when bleaching mainly occurs from the ground state triplet (Figure 2h). With 592 nm co-illumination (Figure 2h, orange line), the effect on photo-switching fatigue recovery depends on the relative strength of each pathway. Our data indicates that in rsEGFP2 photo- bleaching from the excited triplet state is not the dominant contribution allowing photo-switching fatigue to decrease when 592 nm light is applied.

Furthermore, we used the simulation tool to explore the photo-switching fatigue recovery as a function of the power density for 405, 488 and 592 nm illumination (Figure 2i-j and Supplementary Note 8). In the simulated photo- switching fatigue experiments, the 592 nm illumination time was kept constant at 1 ms and overlapped in time with the 488 nm illumination dose. Our simulations suggest that the triplet state can be depopulated effectively by co- illuminating with moderate doses of 592 nm (∼ kW/cm^2^), with the recovery being more evident at higher 405 nm power densities since a relevant fraction needs to be on-switched to yield enough *C^-^* concentration and accumulate triplet state (Figure 2i). Similarly, the photo-switching fatigue recovery is more pronounced at higher 488 nm excitation power densities. In rsEGFP2 at high 488 nm power densities, the off-switching pulse length can be much shorter than the triplet lifetime (< 0.5 ms for 488 nm power densities > 600 W/cm^2^, top half of Figure 2j). In such conditions, the transient triplet population will have a higher probability of interacting only with 592 nm light generating the most photo-switching fatigue recovery. Importantly, the absorption of 592 nm light will not generate additional triplet via ISC but mainly depopulate the transient [*T_1_*] as shown in the scheme in Figure 2c. Nonetheless, the photo-switching fatigue recovery reaches a plateau at 488 nm light > 600 W/cm2 and co-illumination of 592 nm power densities > 10 kW/cm2 which indicates that the triplet state concentration has reached a steady state condition under the experimental illumination parameters.

### Photo-switching fatigue in other green-emitting RSFPs

We extended our investigation of the fatigue resistance to other RSFPs focusing on green negative photo-switchers widely used in multiplexing^10–13^ and super-resolution microscopy^36,42–44^. We observed a correlation between the off-switching speed of the RSFP and the experienced photo-switching fatigue (Figure 3a and Supplementary Note 9), being slower photo-switchers – in particular, rsEGFP(N205S) and rsGreen1 – more prone to bleaching upon cycling than their faster-switching counterparts – rsGFP2 and rsGreenF, respectively -. Practically, a less efficient off-switching process results in longer illumination times per cycle increasing the probability of triplet state formation. Furthermore, if such a state presents a prominent absorption peak at the excitation wavelength, such excess energy will trigger the triplet state excitation and contribute to an enhancement of photo-bleaching.

**Figure 3.**
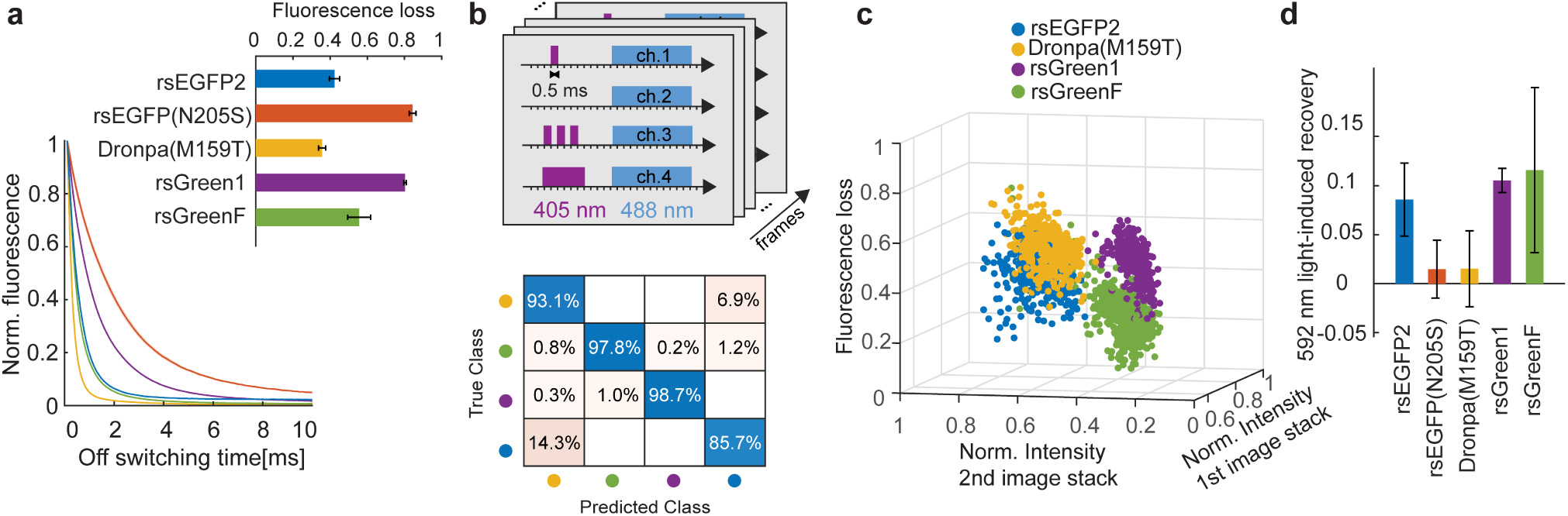
Photo-switching fatigue for multiple green-emitting rsFPs. (a) Off-switching curves for different green negative photo-switchers. The curves were recorded at 405 nm power density 240 W/cm^2^ and 1 ms illumination time; 488 nm power density 200 W/cm^2^. Fluorescence loss due to photo-bleaching for different green negative photo-switchers. The on-switching illumination dose for all the illustrated RSFPs was 1 ms 405 nm at 240 W/cm^2^, while the 488 nm off-switching doses were: 0.9 ms at 300 W/cm^2^ for rsEGFP2; 3.7 ms at 200 W/cm^2^ for rsEGFP(N205S); 0.6 ms at 220 W/cm^2^ for Dronpa(M159T); 1.5 ms at 190 W/cm^2^ for rsGreen1 and 1.0 ms at 240 W/cm^2^ for rsGreenF. (b) On the top, pulse scheme for multiplexing. On every pixel, the sample was illuminated with four 405 nm doses as indicated. To acquire an unmixing frame, the sample was scanned across different multifoci with 90 nm steps resulting in 4 images where the intensity of the features is modulated by the 405 nm dose. The procedure was repeated 15 times to acquire a total of 60 frames. The intensity of every 4^th^ was used to normalize the intensity of the three previous ones. The 4-image intensity traces were averaged to create a 3-dimensional unmixing space. The decay of intensity due to photo-bleaching was also computed as an additional 4^th^ dimension in the unmixing space. On the bottom, the confusion matrix shows the accuracy of the unmixing prediction for 4 RSFPs. The centroids of individual proteins were identified and the whole dataset was reanalyzed using a Gaussian mixture model containing the parameters of isolated clusters. (c) Distribution of segmented bacteria, each expressing a single RSFP, in the 3-dimensional space. This information together with the normalized intensity of the 3^rd^ image stack was used to identify the individual bacteria. (d) Recovery effect for different green negative photo-switchers, when the 592 nm follow the 488 nm pulse delayed of 1 ms.

The distinct fatigue dynamics of each RSFP provide an additional dimension for multiplexing within a single spectral channel^10–13^. These strategies distinguish RSFPs that emit in the same spectral range by analyzing fluorescence signal changes under specific illumination sequences. Methods like LIGHTNING^11^ and TMI^12^ resolve the kinetic rates of light-induced transitions, while approaches such as exNEEMO^10,13^ capture fluorescence intensity changes in response to varying photo-activation sequences. Though versatile and compatible with diverse microscopy set-ups, exNEEMO’s performance declines at high-speed imaging (e.g., confocal imaging with excitation densities ∼ 100 – 500 W cm^-2^) due to overlapping off-switching curves (see Supplementary Note 10).

To address this, we tested photo-bleaching dynamics as an additional distinguishing feature. We designed a pulse scheme with 4 acquisitions per pixel, varying the on-switching energy dose to produce distinct fluorescence responses for each of the 4 proteins (Figure 3b). At each pixel, the sample undergoes the pulse sequence in Figure 3b, with the 4^th^ frame normalizing the recorded intensity of the first three, similar to exNEEMO^10,13^. To account for photo-bleaching, 60 frames – 15 unmixing images – were collected for differently labelled bacteria, and the fluorescence loss was calculated. Data for each RSFP were fitted to a Gaussian model in a 4-dimensional space (Figure 3c), including normalized intensities from the first three image stacks and fluorescence loss. To evaluate the unmixing method, all identified bacteria were reanalyzed using a Gaussian mixture model containing the information of the individual proteins. This pulse scheme is compatible with a parallelized confocal microscope allowing the acquisition of unmixing images with ∼ 0.5 Hz temporal resolution and over extended FOV ∼ 40×40 μm^2^, distinguishing four RSFPs with high accuracy. This approach demonstrates that tracking the photo-switching fatigue in samples with multiple RSFPs enables accurate identification of individual markers.

We then investigate if the reduction of fatigue by red-light co-illumination is a general phenomenon across various RSFPs variants. Our data shows that rsGreen1 and rsGreenF had a significant recovery of photo-switching fatigue upon co-illumination with 592 nm light (Figure 3d). We also measured the recovery as a function of the delay between 488 nm and 592 nm illumination doses (see Supplementary Note 9). The magnitude of the recovery decayed as the pulses were shifted apart suggesting that the effect originates from the depopulation of a ms-lived state, similarly to rsEGFP2. Photo-bleaching pathways linked to the triplet state are compatible with the observed increased photo-switching fatigue in slower photo-switchers. In fact, the slower switching variant rsEGFP(N205S) is not significantly affected by 592 nm co-illumination. Whether the accumulated triplet population may be recovered or not by the red-shifted co-illumination would be controlled by the absorption probability of the triplet and singlet at the red-shifted wavelength and the balance of triplet state quantum yields. Finally, we measured the photo-switching fatigue recovery due to thermal relaxation, which is reduced in slower photo-switchers (see Supplementary Note 9), suggesting that the potential recovered fraction is lower in comparison to faster photo- switchers.

### Prolonged time-lapse imaging at high spatiotemporal resolution with RSFPs

Motivated by our findings in a controlled environment such as a PAA gel or bacteria expressing the different rsFPs, we investigated the strength of the phenomenon in living cells to achieve prolonged time-lapse imaging in different modalities.

Taking advantage of our parallelized confocal microscope^35,45^ we imaged the mitochondrial outer-membrane- protein-25 (OMP-25) labelled with rsEGFP2 in U2OS cells. In such configuration, a spatially patterned 405 nm illumination created by a micro-lens array (MLA) will on-switch fluorophores at determined positions in the sample while the fluorescence is interrogated with another MLA placed in the 488 nm illumination path that is co-aligned to the 405 nm MLA. To form an image, the sample is scanned in between consecutive micro-lens with a step size tuned depending on the target resolution. To achieve a confocal spatial resolution, ∼10 frames – and therefore, photo-switching cycles – are necessary according to the Nyquist sampling criteria. We introduced the 592 nm co- illumination simultaneously with the fluorescence read-out step, aligning a 592 nm illumination with MLA to be more efficient in delivering a sufficient light dose to trigger the optically-induced depopulation. Our data shows a decrease in photo-bleaching after 500 frames with ∼ 10 kW cm^-2^ of 592 nm co-illumination (Figure 4a). As shown in Figure 4a, the features of the mitochondrial network are better preserved after hundreds of imaging frames when the 592 nm dose is added. Moreover, we examined the bleaching constant of individual images in the dataset sorted by their initial signal-to-background ratio – mostly dependent on the transfection efficiency – and observed that for cells displaying similar brightness, the bleaching is slower when the red-shifted co-illumination is added (see Supplementary Note 11). Attaining these results, optically-induced depopulation of the triplet state can prolong time-lapse optical imaging of living cells of dim samples.

**Figure 4.**
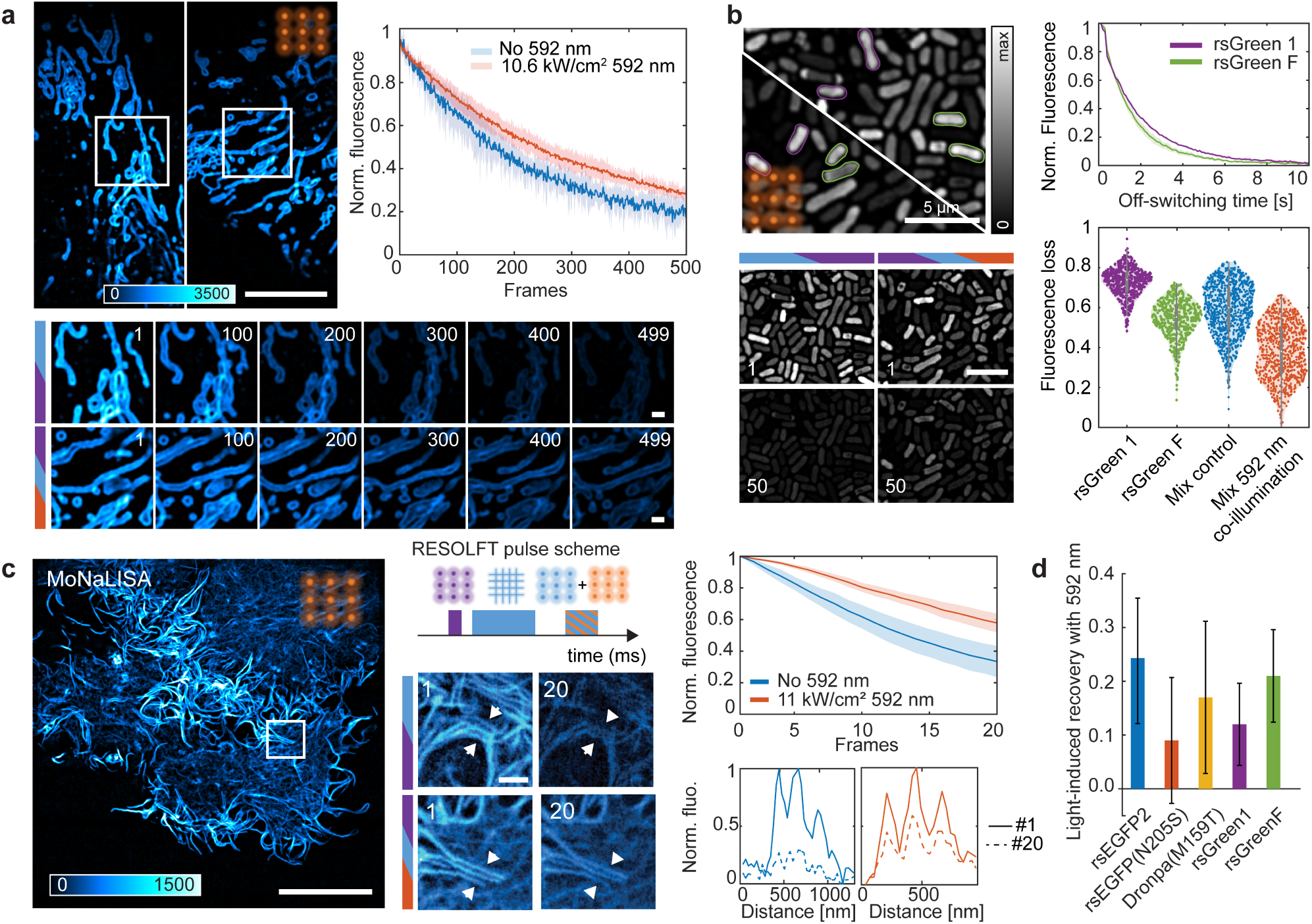
Minimizing photo-switching fatigue in advanced live cell imaging techniques. (a) Fluorescence signal evolution in OMP-25-rsEGFP2 U2OS cells with and without 592 nm illumination imaged in parallelized confocal modality. Each curve is built by averaging bleaching curves of multiple cells (N = 4-5) for each illumination condition. The error was computed as the standard deviation from the mean (± σ) and is represented by the shaded area in each curve. Scale bars, 10 μm and 3 μm. (b) Bacteria sample expressing a mix of rsGreen1 and rsGreenF. The identity of the bacteria is extracted by fitting the off-switching kinetics at the beginning of a confocal timelapse recording of 50 frames, rsGreen 1 as green and rsGreen F as magenta. As the 592 nm is introduced, the fluorescence loss observed for both proteins in the mixed sample decreases and all the bacteria are still identifiable. The violin plot reports on the distribution of fluorescence loss in four conditions: only rsGreen 1, only rsGreen F, mixed sample and mixed sample under 592 nm co-illumination. Scale bars, 5 μm. (c) MoNaLISA imaging for reduced photo-bleaching. The 592 nm light dose is delivered simultaneously with the 488 nm read-out. An example of bleaching reduction in LifeAct-rsEFGP2 U2OS cells is shown. Actin filaments in a cell with 592 nm co-illumination were still visible after 20 frames, while in a cell without 592 nm illumination, these kinds of detailed features are lost after imaging for 20 frames. The graph reports the fluorescence signal evolution in LifeAct-rsEGFP2 U2OS cells with and without 592 nm illumination. Each curve is built by averaging bleaching curves of multiple cells (N = 4-5) for each illumination condition. The error was computed as the standard deviation from the mean (± σ) and is represented by the shaded area in each curve. Scale bars, 10 and 1 μm (d) Light-induced recovery after 20 frames upon 592 nm co-illumination in U2OS cells expressing LifeAct tagged to different green negative photo-switchable fluorescent proteins. The 592 nm power densities used for the different proteins were: 11 kW/cm^2^ for rsEGFP2; 4.8 kW/cm^2^ for rsEGFP(N205S), rsGreen1, rsGreenF; and 6.5 kW/cm^2^ for Dronpa(M159T).

To explore the benefit of red co-illumination in multiplexing imaging we imaged bacteria expressing two RSFPs in confocal modality, rsGreen1 and rsGreenF. The nature of the protein can be identified by resolving the off- switching decay of the proteins like in TMI^12^ (Figure 4b) and the global fluorescence is monitored over a timelapse of 50 frames. Without red co-illumination, the weaker RSFPs set the absolute number of frames of acquisition, while the addition of the red-shifted co-illumination extends the absolute observation window for the multiplexing imaging (Figure 4b). Our data shows that the 592 nm co-illumination can be a powerful aid to extend in time high- resolution single-channel multiplexed imaging with RSFPs in combination with existing unmixing methods.

We further tested the feasibility of the method for super-resolution imaging recordings, specifically in parallelized RESOLFT nanoscopy^35,45^. The spatial confinement necessary is achieved by off-switching the majority of RSFPs located in the periphery of multiple focal spots with a grid-like pattern generated by interfering waves. The on- switching at 405 nm and the fluorescence read-out at 488 nm is carried out with a multi-foci pattern generated with two separate micro-lens arrays (MLAs), which minimize lateral and axial cross-talk.

The parallelization of the imaging scheme minimizes the illumination intensities without compromising the overall frame time. As with any other RESOLFT and NL-SIM imaging scheme, the photo-switching fatigue represents the ultimate limit to the maximum number of frames, which can be recorded.

Similarly to the parallelized confocal imaging experiments, the 592 nm illumination dose is delivered simultaneously and co-aligned to the fluorescence read-out step (Figure 4c). As the majority of the triplet population will reside in the centre of each focal spot, the optically-induced depopulation will be more effective in this position. We observed a decrease in the bleaching fraction after 20 frames in U2OS cells endogenously expressing vimentin- rsEGFP2 with an increasing 592 nm illumination dose (see Supplementary Note 11) as well as in U2OS cells expressing LifeAct-rsEGFP2. Using the same pipeline, we observed a ∼20% reduction in the bleaching fraction after 20 frames (Figure 4c) by adding the red-shifted co-illumination, which competes with reported chemically-based strategies for reducing photo-bleaching^25,26^. As shown in Figure 4c, co-illumination with 592 nm helped in maintaining the signal-to-noise ratio (SNR) during the time-lapses and detailed features of the actin network of the cells were still visible after 20 frames. Similarly, we transfected U2OS cells with LifeAct tagged to other green negative photo-switchers and evaluated the effect of the 592 nm co-illumination in the same manner (Figure 4d). Our data in cells correlates well with our findings in the PAA gel for the different RSFPs showing that this can be a general strategy for reducing photo-switching fatigue in other reversibly switchable fluorescent proteins.

Additionally, we monitored whether the 592 nm co-illumination induced any photo-toxicity in the cells. An assay using the DNA-repairing protein XRCC1 (X-Ray cross-complementary factor 1)^46^, did not report additional DNA photo-damage from the 592 nm illumination dose in the cells (see Supplementary Note 12). Similarly, the mobility of mitochondria was assessed as a photo-toxicity parameter. Both datasets, with and without 592 nm co-illumination, were inspected looking for spontaneous events – such as branching, fission, fusion, long stretching and long retraction – and a time stamp of their occurrence was annotated. Our data shows the similarity in the cumulative distribution of events with and without 592 nm co-illumination (see Supplementary Note 12) suggesting no adverse effects arise from the additional red-shifted illumination regarding mitochondria mobility.

## Discussion

We reported an all-optical solution for maximizing the number of photo-switching cycles in reversibly switchable fluorescent proteins. By studying in depth the loss of fluorescence signal upon photo-switching we identified two different sources: an initial drop as a result of the kinetics of the on-switching process and a more progressive decrease consequence of photo-bleaching. We reported that the former can be mitigated by modifying the delivery of the on-switching energy dose, with an enhanced fluorescence output for longer illumination times and reduced power density. This strategy is compatible with any microscopy modality since it only requires increasing the dwell time.

We proposed a second strategy, based on red-shifted light co-illumination, which mainly benefits the ON-state population of RSFPs. This strategy is particularly suited for microscopy modalities using repetitive light exposure and fast frame rate such as parallelized confocal microscopy, NL-SIM and parallelized RESOLFT. Co-illuminating the sample with a red-shifted wavelength with respect to the fluorescence excitation facilitated the depopulation of a dark state involved in the bleaching pathways of rsEGFP2. We characterized this effect with both experiments and simulations and, based on that knowledge, we proposed a novel pulse scheme for confocal and RESOLFT nanoscopy that effectively mitigates photo-bleaching. Since the phenomenon is seemingly conserved among other green negative photo-switchers, we provide a new imaging solution which advances image multiplexing via RSFPs kinetics. In these experiments, the choice of illumination typically favours one specific variant, for example, a fast switcher, at the cost of higher fatigue for the slower switcher or vice versa. By interleaving red-light, it is possible to uniformize the fatigue curve for different RSFPs variants resulting in higher imaging contrast.

We showed that with the proposed imaging scheme the SNR of fine sub-cellular structures is maintained across tenths of imaging frames. In addition to that, we performed a thorough characterization of the photo-switching fatigue of rsEGFP2 and aided by kinetics simulations we identified two main photo-bleaching pathways, one deriving from the triplet state and another one stemming from the on-switching mechanism of rsEGFP2 that became dominant at high 405 nm illumination doses.

There is a trade-off in any live-cell imaging application between spatiotemporal resolution and photo-stability. In that sense, all-optical approaches to reduce photo-bleaching offer advantages as they do not interfere with optimized protocols in sample preparation and, can be complementary to additional chemically based strategies designed to further improve photo-stability. Our results showed that a deeper understanding of the species involved in the photo- cycle of RSFPs as well as the mechanisms governing photo-bleaching can lead to longer and higher quality live-cell imaging experiments. Moreover, considering the recent spectroscopic characterization of rsEGFP2 at cryo- temperatures^28^, it is reasonable to assume that the photobleaching recovery we report here could be enhanced even further by tailoring the red-shifted co-illumination to the absorption spectrum of the triplet state of rsEGFP2, which is reported to peak around 900 nm^28^. Near-infrared co-illumination successfully extended the number of frames accessible to widefield imaging with EGFP without additional photo-damage^29^. In that regard, the finding of a light-induced depopulation of a dark state mechanism in RSFPs would be impactful in all the applications that utilize photo-switching within biological imaging. From super-resolution imaging approaches such as RESOLFT – as presented here – and pcSOFI, to single protein tracking applications or novel spectroscopy techniques that have photo-switching of fluorescent proteins at their core.

## Methods

### RSFP characterization in PAA gel

All RSFP characterization experiments were done by embedding the protein of interest in polyacrylamide (PAA) gel. To prepare a sample, a few μl (depending on the RSFP) of purified protein were diluted in phosphate buffer for a total volume of 42.0 μl. The protein solution was mixed with 30.0 μl acrylamide (Rotiphorese Gel 30, Roth, Karlsruhe, Germany), 0.75 μl 10% ammonium persulfate (Ammonium peroxydisulfate ≥ 98%, Roth, Karlsruhe, Germany) and 1 μl 10% TEMED (15524-010, Invitrogen). The final solution was thoroughly mixed by vortex and a small volume (10-20 μl) was placed on a glass slide (No 1.5) and a coverslip was pressed onto the sample to produce a thin layer. After complete polymerization, the sample was sealed with silicon-based glue (Picodent twinsil, Picodent, Wipperfürth, Germany) to prevent drying out.

Before each photo-switching fatigue experiment, a characterization of the off and on-switching kinetics of the sample was performed. The off-switching time parametrization was done by averaging 25 off-switching profiles recorded in a small FOV (∼ 1}10 μm^2^) at different 488 nm illumination doses (∼ 10-1000 W/cm^2^). For a given 488 nm power density, the characteristic off-switching time is such that the signal has decayed to 20% of the initial value. To characterize the on-switching response, a 20×20 μm^2^ FOV is consecutively photo-switched with increasing 405 nm illumination doses (∼ 10-1000 W/cm^2^). At each level of 405 nm light, the camera is exposed for the calibrated 488 nm off-switching time and the measurement is repeated 9 times. The mean of the integrated fluorescence for each cycle with its standard deviation is measured and plotted against the 405 nm power density.

To record a photo-switching fatigue experiment, a 20×20 μm^2^ FOV underwent multiple photo-switching cycles (∼ 1600-2200 depending on the experiment). The camera exposure time was matched to the 488 nm pulse width and the integrated fluorescence signal was collected for that photo-switching cycle. At the onset of the new cycle, a 1 ms dose of 405 nm light reset the majority of the ensemble population back to the emissive state. Additionally, a 592 nm wavelength was incorporated into the pulse scheme. The typical pulse dwell time was 10 ms but was adjusted accordingly depending on the 488 nm power density.

The nonlinearity of the on-switching process in rsEGFP2 was tested by modifying the delivery of the 405 nm illumination dose with different combinations of power density and illumination times while keeping a constant total energy of 90 mJ/cm^2^. The total dose of 90 mJ/cm^2^ was chosen by previously calibrating the sample to an on- switching ramp with 1 ms 405 nm illumination time. The same area of 20×20 μm^2^ was repeatedly exposed to 25 on/off photo-switching cycles for every combination of 405 nm power densities (∼ 5–300 W/cm^2^) and illumination times (20-0.3 ms). The fluorescence signal was captured by the camera during the 488 nm illumination. For a given experiment, the FOV is initially illuminated with only 488 nm, the fluorescence retrieved in this initial cycle is considered to represent the true concentration of fluorophores in that volume. The fluorescence output for every on-switching condition is normalised to the integrated signal in that initial photo-switching cycle for every FOV. Each data point was built by averaging 20 of the 25 photo-switching cycles for a given condition. The experiment was repeated in 4 different areas of a rsEGFP2-embedded PAA gel.

Moreover, to characterize processes with a higher temporal resolution a confocal system equipped with an avalanche photodiode (APD) detection system was used. To examine the on-switching mechanism, a burst (5 μs) of high-power 405 nm light (∼ 30 kW/cm^2^) triggered the on-switching transition, after a modulable delay (5 μs – 50 ms), the subsequent fluorescence signal was read out with 488 nm illumination at ∼ 40 kW/cm^2^. The area below the off-switching curve was calculated and plotted against the variable delay. Similarly, we investigated the relaxation of a dark state formed upon 488 nm absorption from the emissive state. In this case, a 5 μs 405 nm burst (∼ 30 kW/cm^2^) was used as the on-switching light dose, after a 3 ms delay – to ensure full relaxation to the emissive state – a first 488 nm dose (500 μs at ∼ 40 kW/cm^2^) was delivered to the sample, enough to decrease the observed signal down to 80-90 % of the initial value. After that, a second 488 nm pulse with the same illumination dose was delivered. The delay between consecutive 488 nm pulses systematically varied between 5 μs and ∼ 500 ms. The area below the second off-switching curve was compared to a control curve – where no delay existed between pulses – and plotted against the delay time. All the experimental parameters are reported in Supplementary Table 7.

### Optical set-ups for characterization experiments

The photo-switching fatigue experiments, as well as the characterization experiments, were carried out on a modified version of the set-up reported here^45^ (see Supplementary Figure 20). In brief, 3 different wavelengths were combined in a WF microscope configuration with a high numerical aperture (NA) objective (HCX PC APO 100×/1.40–0.70-NA oil-immersion; Leica Microsystems). Photo-activation and fluorescence excitation were carried out using modulable laser diodes, 405 nm (200 mW, 06- 01, 405 nm; Cobolt) and 488 nm (200 mW, 06-01, 488 nm; Cobolt) respectively. To test the photo-switching fatigue recovery, three red-shifted wavelengths were incorporated, 592 nm (DEMO-2RU-VFL-P-2000-592; MPB Communications) modulated by an acousto-optic tunable filter (AOTF, AOTF-nC-VIS-TN; AA Optoelectronics). The emitted fluorescence was directed to the detection path with a dichroic mirror (ZT405/488//640/775RDC; Chroma) and later to a scientific CMOS (sCMOS) camera (ORCA Fusion C14440-20UP; Hamamatsu). In the microscope, the light doses were digitally modulated by transistor-transistor logic signals with a NIDAQ PCI 6371 (National Instrument) acquisition device. The NIDAQ instrument synchronized the pulse sequence and the camera readout. The length and delay of the laser pulses, the output power of the lasers, the exposure time of the camera, and all other relevant parameters can be controlled through ImSwitch^47^. For sample screening, we added a wide- field path, which we could alternate with the main WF illumination by flipping a motorized mirror. The power densities of the different beams were calculated by measuring the power at the objective back-aperture and the area was computed by fitting a Gaussian function to the beam profile imaged on an AlexaFluor488 thin layer, for 405 and 488 nm beams, and Abberior STAR RED, for 592 nm beam.

Experiments where higher temporal resolution was required were performed using the confocal microscope configuration shown here**^8^**(see Supplementary Figure 21). In brief, 405 and 488 nm lasers were combined and tightly focused to a diffraction-limited volume with a high NA objective (APO 100×/ 1.40 NA oil immersion; Leica Microsystems). Photo-activation and fluorescence excitation were carried out using modulable laser diodes, 405 nm (125 mW, 06-01, 405 nm; Cobolt) and 488 nm (70 mW, 06-01, 488 nm; Cobolt) respectively. The 405 nm polarization was controlled with achromatic λ/2 and λ/4 mounted on remotely controlled rotating stages (Thorlabs) and set to circular polarization. To record an experiment, the microscope incorporates a *xy* galvo scanning system (Cambridge Technology) to scan the light beams across the FOV. The emitted fluorescence is later de-scanned and directed to a pair of APDs (MPD). The counts recorded in both detectors are summed together later in the data treatment. The microscope components were controlled using a field-programmable gate array card (FPGA, PCIe- 7852, National Instruments) and custom-designed software in LabVIEW. The power densities of the different beams were calculated by measuring the power at the objective back-aperture and the area was computed by calculating the FWHM of the beam’s PSF from a gold beads sample. The schematic of the microscopes as well as a list of the optical components are reported in Supplementary Note 13.

The experiments characterizing the photo-switching fatigue recovery effect of near-infrared (NIR) triplet absorption on rsEGFP2 photo-switching fatigue were performed on an epi-illuminated, commercial confocal laser scanning microscope (Olympus FV1200). The setup was integrated by a combination of elements from two previous works^48,49^ (see Supplementary Figure 22). The PAA-embedded rsEGFP2 was on-switched by a train of 405 nm (LDH-D-C-405, PicoQuant GmbH, Berlin) picosecond pulses extending for 1 ms at 80 MHz repetition rate and an average power density of ∼ 300 W/cm^2^ at the sample plane. After a 1 ms delay, the confocal volume was illuminated with a train of picosecond 488 nm (LDH-D-C-485, PicoQuant GmbH, Berlin) pulses spanning 1 ms at the same repetition rate as the 405 nm laser and an average power density of ∼ 14 kW/cm^2^ at the sample plane. The diode lasers were controlled via a laser driver (Sepia PDL 828) with a repetition rate of 80 MHz. Since the pulse width of the laser was less than 100 ps, a train of pulses constituted a larger pulse. The pulse trains obtained for the final measurements are as mentioned along with the optical setup schematic in Supplementary Note 13. After a 10 ms period of no illumination, the photo-switching cycle was repeated again for ∼ 5000 cycles.

The NIR co-illumination was provided by a tunable Ti: Sapphire laser (Mira 900, Coherent, pumped by Nd: Vanadate laser (Verdi™ V-10, Coherent)) and was tuned to 810 nm and 900 nm for scanning the triplet spectrum. It was operated in a continuous mode and it’s input was controlled using a mechanical shutter. Contrary to the characterization experiments in the WF configuration, the NIR co-illumination was applied during the whole duration of the photo-switching fatigue experiment. The power of the Ti: Sapphire laser was varied to achieve power densities of ∼ 1- 50 kW/cm^2^ at the sample plane for both wavelengths (810 and 900 nm). The fluorescence was collected back from the same objective (UPlanSApo 60×/1.2w, Olympus) and then passed through a dichroic mirror (ZT405/488/635rpc-UF2, Chroma) after which the light was focused onto a pinhole of 100 μm. A slightly wider pinhole size was chosen in order to have a larger volume to average deformities present at every focal plane of the gel. A bandpass filter (535/70) was used in the emission path and finally, the light was collected by two avalanche photodiodes (Tau-SPAD, PicoQuant GmbH, Berlin). The signals were directed to a data acquisition system (Hydraharp 400, Picoquant, Berlin) and time traces of the signals were exported via Symphotime and analysed with a home-built code.

### rsEGFP2 photo-physical modelling

To provide an insight on the mechanisms involved in the photo-switching fatigue we built a photo-physics simulation tool. To simulate the response of rsEGFP2, we emulated the experimental framework and calculated the population evolution given a proposed kinetic scheme. The system of kinetic equations is solved in matrix form. Since the photophysical evolution of the RSFP chromophore can be specified as a linear system of ordinary differential equations, it is possible to find an analytical solution to propagate a set of initial populations for each species. The solution involves the diagonalization of the kinetic rate matrix^50^, which collects all the rates connecting the different species among each other. In general, to simulate a complex pulse scheme we divide it into time windows – instances in which the laser states are not changing and thus the kinetic rates are fixed – and solve the rate equations for the length of such window. For several consecutive time windows, the populations of the species at the last simulation point of one window are the initial populations of the next window. A more detailed description of the theory behind the photo-physical simulations can be found in Supplementary Note 14.

The proposed photo-cycle model includes the 4 main states responsible for the photo-switching in rsEGFP2 – *TH, CH, C^-^, T^-^* - and an intermediate species, as described in the literature^15^. Furthermore, to parametrize the observed photo-switching fatigue we included two bleaching pathways: (i), triplet states bleaching, and (ii), photo-destruction of the chromophore from the off-state. The complete photo-cycle scheme with the different transitions between states is shown in Supplementary Note 1. Attaining the different levels of complexity that the photo-cycle model includes, different pathways were activated or silenced depending on the scope of the simulation. The parameters input in the model are taken from literature, properly referenced, or by fitting the model to our data. The parameters used for each of the reported simulations can be found in Supplementary Table 2.

### Protein purification

Transformed E. coli cells (TOP10 and BL21 strain, pQEpBAD, and pQE expression system) were grown overnight in LB medium supplemented with ampicillin. The next morning, cells were diluted 1:100 together with ampicillin 1:1000 in LB medium and added to 50-ml tubes so that the tubes were filled to the brim, and the tubes were then sealed and incubated until the optical density (OD) at 600 nm reached 0.6 at 37 °C. Then IPTG was added to a final concentration of 1 mM, and the tubes were tightly sealed again to restrict oxygen availability. After overnight incubation, the cells were collected by centrifugation in the same tubes. The resulting bacterial pellet was sonicated and re-concentrated by centrifugation. The remaining supernatant was filtered (MinSart RC Hydrophilic 25 mm 0.2 μm, Sartorius, Göttingen, Germany) and the proteins were purified by Ni-NTA affinity chromatography (His SpinTrap; GE Healthcare) according to the manufacturer’s instructions. The purified proteins were concentrated by ultrafiltration and taken up in phosphate buffer, pH 7.5.

### RESOLFT and parallelized confocal microscopy

Live-cell RESOLFT and parallelized confocal imaging were performed on a modified version of the microscope set-up reported here^45^, the same one used for the photo-physical characterization at low illumination power densities (see Supplementary Figure 20). To achieve spatial resolutions below the diffraction limit, the light doses were spatially patterned in the sample plane by introducing optical elements in the laser paths. Photo-activation was carried out at 405 nm in combination with a micro-lens array (MLA, MLA-150-5C-M; Thorlabs) to create a multi-foci pattern with 625 nm periodicity. Illumination at 488 nm and 592 nm were combined using a dichroic mirror (zt502rdc; Chroma) and focused by an MLA to create a multi-foci pattern at the sample plane with periodicity 625 nm and that was aligned to the 405 nm illumination. To create the confinement light pattern, the collimated light of the 488 nm digitally modulated diode laser (200 mW, 06-01, 488 nm; Cobolt) was directed to a half-wave plate and polarizing beam splitter (PBS). The half-wave plate was adjusted to equally split the light according to the p and s orientations. After the polarizing beam splitter, the light was sent through custom-made phase-diffraction gratings of 437-nm-high SiO2 lines with a 25 μm period (Laser Laboratorium Göttingen). After the gratings, the light paths were recombined with another polarizing beam splitter. The + 1 and –1 diffraction orders at the back focal plane of the objective to obtain a final OFF periodicity of 312.5 nm. A piezo scanning system (BPC303; Thorlabs) was used to apply pulse illuminations across a small region equal to the multi-foci periodicity at precise steps (typically 35 nm but adjusted to the desired optical resolution). The light doses were digitally modulated by transistor-transistor logic signals with a NIDAQ PCI 6371 (National Instrument) acquisition device. The FOV for MoNaLISA imaging was ∼ 40×40 μm^2^. The typical imaging pulse scheme consisted of 4 illumination sequences for RESOLFT imaging – on-switching, confinement, read-out and recovery, and 3 illumination sequences for parallelized confocal imaging – on-switching, read-out and recovery. The NIDAQ instrument synchronized the pulse sequence, the camera readout, and the scanning. The readout laser was synchronized with the sCMOS (ORCA Fusion C14440-20UP; Hamamatsu) camera exposure to detect fluorescent emission. The length and delay of the laser pulses, the output power of the lasers, the exposure time of the camera, and all other relevant parameters were controlled through ImSwitch^47^. A schematic of the imaging microscope as well as a list of the optical components can be found in Supplementary Note 13. The imaging parameters are reported in Supplementary Table 8.

### Cell culture and transfection

Wild-type U2OS (ATCC HTB-96) cells and U2OS with endogenously tagged vimentin-rsEGFP2^51^ were cultured in DMEM (Thermo Fisher Scientific; 41966029) supplemented with 10% (vol/vol) FBS (Thermo Fisher Scientific; 10270106) and 1% penicillin-streptomycin (Sigma Aldrich; P4333) and maintained at 37 °C and 5% CO2 in a humidified incubator. For transfection, 2 ×10^5^ cells per well were seeded on coverslips in a six-well plate. After 1 d, cells were transfected using Lipofectamine LTX Reagent with PLUS reagent (Thermo Fisher Scientific; 15338100) according to the manufacturer’s instructions. 24–36 h after transfection, cells were washed in PBS solution, placed with phenol-red-free DMEM or Leibovitz’s L-15 medium (Thermo Fisher Scientific; 21083027) in a chamber, and imaged.

### Live-cell single channel multiplexing

Multiplexing experiments were carried out on transformed E. coli cells prepared as mentioned above. Bacteria expressing single RSFPs were imaged after centrifugation and mixed samples were prepared by combining bacteria expressing different proteins at suitable concentrations. Two different methods for unmixing were employed: recording of slow off-switching curves and intra-pixel 405 nm intensity modulation inspired by exNEEMO^10,13^. For the former, the bacteria in the FOV were off-switched using the 488 nm illumination (∼ 5 μW at the sample plane) of the auxiliary WF shown in Supplementary Figure 20 with ∼ 100 ms exposure time. Afterwards, the same bacteria were imaged in the parallelized confocal modality in the microscope described above. The scanning step between adjacent multi-foci was set to 90 nm. Since the WF and parallelized confocal images have different coordinates systems, the image stacks were transformed and registered with the MultiStackReg^52^ plug-in in ImageJ. Following the registration, a custom-written segmentation routine in ImageJ was used to identify single bacteria in both WF and parallelized confocal image stacks and the intensity traces were extracted. The data clustering is performed by a custom-written Matlab script based on a Gaussian mixture model (GMM) that fits the data into a given number of Gaussian components, extracting μ and σ.

On the other hand, the intra-pixel 405 nm intensity modulation method consisted of a series of 4-frame parallelized confocal stacks with modulated 405 nm illumination doses. On every pixel, the sample is consecutively illuminated with 4 different pulse sequences the integrated intensity is collected during the 488 nm excitation and the modulation of the fluorescence levels is given by the on-switching dose. The sample is scanned between adjacent multi-foci with a step of 90 nm. The microscope control software ImSwitch^47^ was adapted to acquire N-images per pixel. A custom-written Python script splits the acquired frames into 4 and our reconstruction algorithm returns a stack of 4 frames, each with a modulated pixel intensity given by the input illumination sequence, for every acquisition frame. To build a time-lapse recording, this procedure is repeated 15 times resulting in a total of 60 frames. Following the acquisition, a custom-written script in ImageJ is used for image processing and segmentation of individual bacteria. The intensity traces of the identified bacteria are extracted and processed in a custom-written Matlab script. Similarly to exNEEMO^10,13^, the intensity of every 4^th^ frame is used to normalize the intensity of the three previous ones and the average of the normalized 4-frame stack intensity trace across the 60 frames time-lapse is computed. Simultaneously, the fluorescence signal loss due to bleaching is computed as the difference in average raw intensity between the first 3 normalization frames and the last 3 normalization frames in the stack. The data clustering is performed by a custom-written Matlab script based on a Gaussian mixture model (GMM) that fits the data into a given number of Gaussian components. The first training step extracts μ and σ for the individual proteins, afterwards the whole dataset is reanalyzed using the fitted Gaussian mixture model and the confusion matrix is created.

## Data availability

The data Supplementary the findings of this study are available from the corresponding authors upon request.

## Code availability

The code for the photophysics simulation of rsFPs is available at the GitHub repository https://github.com/marinaguilera/photophysics_simulator.git.

## Corresponding Author

Francesca Pennacchietti – SciLifeLab and KTH Royal Institute of Technology, 17165, Solna, Sweden; Email:

Ilaria Testa – SciLifeLab and KTH Royal Institute of Technology, 17165, Solna, Sweden; Email:

Author Contributions

I.T. designed the research. I.T. and F.P. co-supervised the research. F.P., A.P. and G.M.A. performed the experiments and analysed the data. A.V. and G.M.A. developed the model. J.W., N.B., A.K., and G.M.A. contributed to the spectral experiments in the near-infrared. G.M. and G.M.A. modified the microscope control software ImSwitch to acquire the multiplexing recordings. I.T., F.P. and G.M.A. wrote the manuscript with the help of all the authors.

## References

1. Hofmann, M., Eggeling, C., Jakobs, S. & Hell, S. W. Breaking the diffraction barrier in fluorescence microscopy at low light intensities by using reversibly photoswitchable proteins. Proc. Natl. Acad. Sci. 102, 17565–17569 (2005).

2. Grotjohann, T. et al. Diffraction-unlimited all-optical imaging and writing with a photochromic GFP. Nature 478, 204–208 (2011).

3. Rego, E. H. et al. Nonlinear structured-illumination microscopy with a photoswitchable protein reveals cellular structures at 50-nm resolution. Proc. Natl. Acad. Sci. 109, (2012).

4. Dedecker, P., Mo, G. C. H., Dertinger, T. & Zhang, J. Widely accessible method for superresolution fluorescence imaging of living systems. Proc. Natl. Acad. Sci. 109, 10909–10914 (2012).

5. Egner, A. et al. Fluorescence Nanoscopy in Whole Cells by Asynchronous Localization of Photoswitching Emitters. Biophys. J. 93, 3285–3290 (2007).

6. Willig, K. I., Stiel, A. C., Brakemann, T., Jakobs, S. & Hell, S. W. Dual-Label STED Nanoscopy of Living Cells Using Photochromism. Nano Lett. 11, 3970–3973 (2011).

7. Gao, Z. et al. Scanning Switch-off Microscopy for Super-Resolution Fluorescence Imaging. Nano Lett. acs.nanolett.4c02452 (2024) doi:10.1021/acs.nanolett.4c02452.

8. Volpato, A. et al. Extending fluorescence anisotropy to large complexes using reversibly switchable proteins. Nat. Biotechnol. 41, 552–559 (2023).

9. Subach, F. V. et al. Red Fluorescent Protein with Reversibly Photoswitchable Absorbance for Photochromic FRET. Chem. Biol. 17, 745–755 (2010).

10. Valenta, H. et al. Separation of spectrally overlapping fluorophores using intra-exposure excitation modulation. Biophys. Rep. 1, 100026 (2021).

11. Chouket, R. et al. Extra kinetic dimensions for label discrimination. Nat. Commun. 13,1482 (2022).

12. Qian, Y., Celiker, O. T., Wang, Z., Guner-Ataman, B. & Boyden, E. S. Temporally multiplexed imaging of dynamic signaling networks in living cells. Cell 186, 5656–5672.e21 (2023).

13. Valenta, H. et al. Per-pixel unmixing of spectrally overlapping fluorophores using intra-exposure excitation modulation. Talanta 269, 125397 (2024).

14. Coquelle, N. et al. Chromophore twisting in the excited state of a photoswitchable fluorescent protein captured by time-resolved serial femtosecond crystallography. Nat. Chem. 10, 31–37 (2018).

15. Woodhouse, J. et al. Photoswitching mechanism of a fluorescent protein revealed by time-resolved crystallography and transient absorption spectroscopy. Nat. Commun. 11, 741 (2020).

16. Uriarte, L. M. et al. Structural Information about the *trans* -to- *cis* Isomerization Mechanism of the Photoswitchable Fluorescent Protein rsEGFP2 Revealed by Multiscale Infrared Transient Absorption. J. Phys. Chem. Lett. 13, 1194–1202 (2022).

17. Habuchi, S. et al. Reversible single-molecule photoswitching in the GFP-like fluorescent protein Dronpa. Proc. Natl. Acad. Sci. 102, 9511–9516 (2005).

18. Yadav, D. et al. Real-Time Monitoring of Chromophore Isomerization and Deprotonation during the Photoactivation of the Fluorescent Protein Dronpa. J. Phys. Chem. B 119, 2404–2414 (2015).

19. Mizuno, H. et al. Higher resolution in localizationmicroscopy by slower switching of a photochromic protein. Photochem. Photobiol. Sci. 9, 239–248 (2010).

20. Zhang, X. et al. Development of a Reversibly Switchable Fluorescent Protein for Super-Resolution Optical Fluctuation Imaging (SOFI). 9, (2015).

21. Moeyaert, B. & Vandenberg, W. SOFIevaluator: a strategy for the quantitative quality assessment of SOFI data.

22. Quérard, J. Resonant out-of-phase fluorescence microscopy and remote imaging overcome spectral limitations. Nat. Commun.

23. Mamontova, A. V., Grigoryev, A. P., Tsarkova, A. S., Lukyanov, K. A. & Bogdanov, A. M. Struggle for photostability: Bleaching mechanisms of fluorescent proteins. Russ. J. Bioorganic Chem. 43, 625– 633 (2017).

24. Greenbaum, L., Rothmann, C., Lavie, R. & Malik, Z. Green Fluorescent Protein Photobleaching: a Model for Protein Damage by Endogenous and Exogenous Singlet Oxygen. Biol. Chem. 381, (2000).

25. Bogdanov, A. M. Cell culture medium affects GFP photostability: a solution. (2009).

26. Roebroek, T., Duwé, S., Vandenberg, W. & Dedecker, P. Reduced Fluorescent Protein Switching Fatigue by Binding-Induced Emissive State Stabilization. Int. J. Mol. Sci. 18, 2015 (2017).

27. Byrdin, M., Duan, C., Bourgeois, D. & Brettel, K. A Long-Lived Triplet State Is the Entrance Gateway to Oxidative Photochemistry in Green Fluorescent Proteins. J. Am. Chem. Soc. 140, 2897– 2905 (2018).

28. Rane, L. et al. Light-Induced Forward and Reverse Intersystem Crossing in Green Fluorescent Proteins at Cryogenic Temperatures. J. Phys. Chem. B 127, 5046–5054 (2023).

29. Ludvikova, L. et al. Near-infrared co-illumination of fluorescent proteins reduces photobleaching and phototoxicity. Nat. Biotechnol. (2023) doi:10.1038/s41587-023-01893-7.

30. Ringemann, C. et al. Enhancing Fluorescence Brightness: Effect of Reverse Intersystem Crossing Studied by Fluorescence Fluctuation Spectroscopy. ChemPhysChem 9, 612–624 (2008).

31. Widengren, J. & Seidel, C. A. M. Manipulation and characterization of photo-induced transient states of Merocyanine 540 by fluorescence correlation spectroscopy. Phys. Chem. Chem. Phys. 2, 3435– 3441 (2000).

32. Richards, C. I., Hsiang, J.-C. & Dickson, R. M. Synchronously Amplified Fluorescence Image Recovery (SAFIRe). J. Phys. Chem. B 114, 660–665 (2010).

33. Hsiang, J.-C., Jablonski, A. E. & Dickson, R. M. Optically Modulated Fluorescence Bioimaging: Visualizing Obscured Fluorophores in High Background. Acc. Chem. Res. 47, 1545–1554 (2014).

34. Grotjohann, T. et al. rsEGFP2 enables fast RESOLFT nanoscopy of living cells. eLife 1, e00248 (2012).

35. Masullo, L. A. et al. Enhanced photon collection enables four dimensional fluorescence nanoscopy of living systems. Nat. Commun. 9, 3281 (2018).

36. Bodén, A. et al. Volumetric live cell imaging with three-dimensional parallelized RESOLFT microscopy. Nat. Biotechnol. 39, 609–618 (2021).

37. El Khatib, M., Martins, A., Bourgeois, D., Colletier, J.-P. & Adam, V. Rational design of ultrastable and reversibly photoswitchable fluorescent proteins for super-resolution imaging of the bacterial periplasm. Sci. Rep. 6, 18459 (2016).

38. Laptenok, S. P. et al. Infrared spectroscopy reveals multi-step multi-timescale photoactivation in the photoconvertible protein archetype dronpa. Nat. Chem. 10, 845–852 (2018).

39. Nienhaus, K. & Nienhaus, G. U. Chromophore photophysics and dynamics in fluorescent proteins of the GFP family. J. Phys. Condens. Matter 28, 443001 (2016).

40. Bourges, A. C. et al. Quantitative determination of the full switching cycle of photochromic fluorescent proteins. Chem. Commun. 59, 8810–8813 (2023).

41. Peng, B. et al. Optically Modulated and Optically Activated Delayed Fluorescent Proteins through Dark State Engineering. J. Phys. Chem. B 125, 5200–5209 (2021).

42. Chmyrov, A. et al. Nanoscopy with more than 100,000 ‘doughnuts’. Nat. Methods 10, 737–740 (2013).

43. Casas Moreno, X., et al. Multi-foci parallelised RESOLFT nanoscopy in an extended field-of-view. J. Microsc. 291, 16–29 (2023).

44. Duwé, S. et al. Expression-Enhanced Fluorescent Proteins Based on Enhanced Green Fluorescent Protein for Super-resolution Microscopy. ACS Nano 9, 9528–9541 (2015).

45. Pennacchietti, F. et al. Fast reversibly photoswitching red fluorescent proteins for live-cell RESOLFT nanoscopy. Nat. Methods 15, 601–604 (2018).

46. Serebrovskaya, E. O. et al. Light-induced blockage of cell division with a chromatin-targeted phototoxic fluorescent protein. Biochem. J. 435, 65–71 (2011).

47. Moreno, X., Al-Kadhimi, S., Alvelid, J., Bodén, A. & Testa, I. ImSwitch: Generalizing microscope control in Python. J. Open Source Softw. 6, 3394 (2021).

48. Sandberg, E. et al. Photoisomerization of Heptamethine Cyanine Dyes Results in Red-Emissive Species: Implications for Near-IR, Single-Molecule, and Super-Resolution Fluorescence Spectroscopy and Imaging. J. Phys. Chem. B 127, 3208–3222 (2023).

49. Bagheri, N., Chen, H., Rabasovic, M. & Widengren, J. Non-fluorescent transient states of tyrosine as a basis for label-free protein conformation and interaction studies. Sci. Rep. 14, 6464 (2024).

50. Berberan-Santos, M. N. & Martinho, J. M. G. The integration of kinetic rate equations by matrix methods. J. Chem. Educ. 67, 375 (1990).

51. Ratz, M., Testa, I., Hell, S. W. & Jakobs, S. CRISPR/Cas9-mediated endogenous protein tagging for RESOLFT super-resolution microscopy of living human cells. Sci. Rep. 5, 9592 (2015).

52. Thevenaz, P., Ruttimann, U. E. & Unser, M. A pyramid approach to subpixel registration based on intensity. IEEE Trans. Image Process. 7, 27–41 (1998).

