## Supporting Information for "All-Optical Strategies to Minimize Photo-Bleaching in Reversibly Switchable Fluorescent Proteins"

**Supplementary Note 1: Modelling of rsEGFP2 photo-cycle**

To assess and interpret the results obtained from the experimental characterization of rsEGFP2, we developed a simulation tool that reproduces the experiment's framework given a photo-switching scheme. Supplementary Figure 1 below shows the complete photo-switching scheme used in this work.

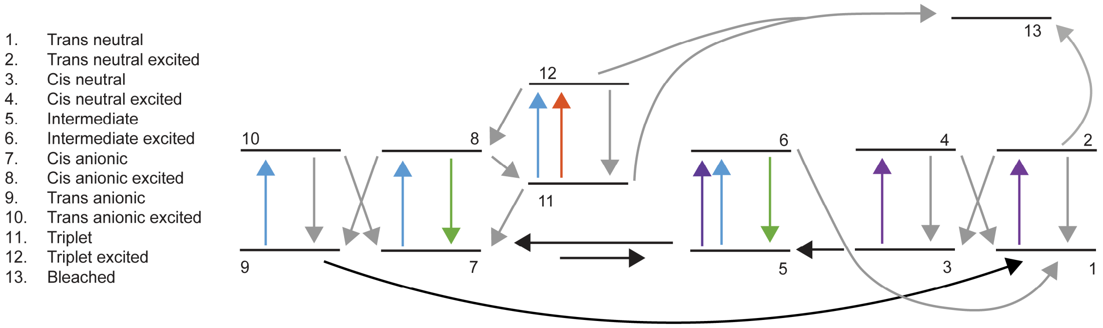


**Supplementary Figure 1. Kinetic scheme for rsEGFP2: 1-Trans neutral; 2-Trans neutral excited; 3- Cis neutral; 4-Cis neutral excited; 5-Intermediate, 6-Intermediate excited; 7- Cis anionic; 8-Cis anionic excited; 9-Trans anionic; 10-Trans anionic excited; 11-Triplet; 12-Triplet excited; 13-Bleached. The coloured arrows (cyan, violet and orange) indicate the wavelength that mainly drives the transition between electronic states. Green arrows indicate fluorescence emission from that state. Grey-coloured arrows indicate a non-radiative relaxation or excited state reaction. Black arrows indicate ground state transitions.**

The scheme above accounts for the excited state isomerization reactions that drive the photo-switching, the presence of the intermediate state (*Int)* in the on-switching process, the protonation/deprotonation processes, the formation of the triplet state from the fluorescent form and the different effective bleaching channels. The cyan, violet, and orange arrows aim to represent the wavelength that mainly drives the transition between the different electronic states. The green arrows represent fluorescent emission, both from the *Int* (6) and the *Cis anionic* ($C^-$) (8) forms. Grey arrows describe processes from the excited state: non-radiative relaxations or excited state reactions. Black arrows depict transitions that occur in the ground state. The different transitions between states are listed in Supplementary Table 1 as i.e. $K_{21}$, which indicates a process that moves population ***to* state *2 from* state *1***.

As presented, the proposed kinetic scheme aims to reproduce three main phenomena: the photo-switching of rsEGFP2 (on-to-off and off-to-on transitions); the formation of triplet state via intersystem crossing and the triplet's interaction with light; and lastly, the observed loss of signal due to photo-bleaching. The purpose of the simulation was to parametrize and help clarify the photo-physical pathways that could describe the experimental observations, therefore, not all the simulated experiments required the same level of complexity from the kinetic scheme. A reduced model including only electronic states 1-10 of Supplementary Figure 1 was used to estimate the absorption parameters of the *Int* as discussed in Supplementary Note 2-3. Since such experiments did not require many photo-switching cycles, the formation of a *triplet* state nor the different bleaching pathways were included in the modelling of rsEGFP2. On the other hand, the full kinetic scheme of Supplementary Figure 1 was required to estimate the absorption properties of the *triplet* state as well as when discussing the different bleaching pathways that intervene in the destruction of the fluorescence signal upon cycling.

Some assumptions were made when building up the scheme. For example, triplet formation via intersystem crossing only appears from the $C^-$ although the *Int* state is also described as fluorescent. We reason that since $\varepsilon^{on}_{488} > \varepsilon^{int}_{488}$, any triplet state population build-up will mostly come from the $C^-$. Regarding the *Int* state as well, the isomerization back to the off-state (transition 6 → 1) has been assumed to be a one-step excited state reaction, however, it is also plausible that the transition is mediated by a cascade of intermediary states with characteristic lifetimes. The direct protonation of the *cis anionic* form of the chromophore (transition 7 → 5) has been accounted for, nonetheless, the importance of such reaction at experimental pH = 7.5 is negligible since the $pK_a^{cis} = 5.9$. On the other hand, the deprotonation of the *trans neutral* (*TH)* isomer (transition 1 → 9, *TH* to *trans anionic,* $T^-$) has not been included as $pK_a^{trans} >> pH$. Moreover, electronic states with the same protonation state have been considered to share the same spectroscopic properties, that is the extinction molar coefficients, $\varepsilon^{CH}_{\lambda} \approx \varepsilon^{TH}_{\lambda}$ and $\varepsilon^{C^-}_{\lambda} \approx \varepsilon^{T^-}_{\lambda}$ as well as the quantum yields for the *cis-trans* isomerization $\Phi_{TH\rightarrow CH} \approx \Phi_{CH\rightarrow TH}$ and $\Phi_{C^-\rightarrow T^-} \approx \Phi_{T^-\rightarrow C^-}$. From Supplementary Table 1, $\sigma(\lambda)$ refers to the absorption cross-section for a given wavelength in each state. $\sigma(\lambda)$ is calculated from $\varepsilon(\lambda)$ as indicated in literature[1]. $\rho(\lambda)$ is the photon flux delivered to the sample as calculated from the power density measured at the objective's back aperture and the size and shape of the beam profile. $\Phi$ refers to the quantum yield of different processes and $\tau$ are the lifetimes of the excited states. All the parameters used in the simulations are listed and referenced below in Supplementary Table 2.

**Supplementary Table 1. Index of the transitions between electronic states of Supplementary Figure 1. The different transitions are described as i.e. K21, which indicates a process that moves the population *to state 2 from state 1*.**

| K | Equation | Description |
|---|---|---|
| $K_{21}$ | $\sigma_{TH}(\lambda) * \rho(\lambda)$ | Trans neutral absorption |
| $K_{12}$ | $1/\tau_{TH}$ | Trans neutral relaxation |
| $K_{32}$ | $\Phi_{TH\rightarrow CH}/\tau_{TH}$ | Trans-to-cis neutral isomerization |
| $K_{43}$ | $\sigma_{CH}(\lambda) * \rho(\lambda)$ | Cis neutral absorption |
| $K_{34}$ | $1/\tau_{CH}$ | Trans neutral relaxation |
| $K_{14}$ | $\Phi_{CH\rightarrow TH}/\tau_{CH}$ | Cis-to-trans neutral isomerization |
| $K_{53}$ | $1/\tau_{Rearrangement}$ | Reorganization of the chromophore in the pocket |
| $K_{65}$ | $\sigma_{Int}(\lambda) * \rho(\lambda)$ | Intermediate absorption |
| $K_{56}$ | $\Phi_{fluo}/\tau_{INT}$ | Fluorescence emission |
| $K_{16}$ | $\Phi_{INT\rightarrow TH}/\tau_{INT}$ | Intermediate-to-trans-neutral isomerization |
| $K_{75}$ | $1/\tau_{Deprotonation-CH}$ | Deprotonation of the chromophore in the cis form |
| $K_{57}$ | $1/\tau_{protonation-C-}$ | Protonation of the chromophore in the cis form |
| $K_{87}$ | $\sigma_{C-}(\lambda) * \rho(\lambda)$ | Cis anionic absorption |
| $K_{78}$ | $\Phi_{fluo}/\tau_{C-}$ | Fluorescence emission |
| $K_{98}$ | $\Phi_{C-\rightarrow T-}/\tau_{C-}$ | Cis-to-trans anionic isomerization |
| $K_{109}$ | $\sigma_{T-}(\lambda) * \rho(\lambda)$ | Trans anionic absorption |
| $K_{910}$ | $1/\tau_{T-}$ | Trans anionic relaxation |
| $K_{710}$ | $\Phi_{T-\rightarrow C-}/\tau_{T-}$ | Trans-to-cis anionic isomerization |
| $K_{19}$ | $1/\tau_{protonation-T-}$ | Protonation of the chromophore in the trans isomer |
| $K_{118}$ | $\Phi_{ISC}/\tau_{C-}$ | Triplet state formation via intersystem crossing |
| $K_{711}$ | $1/\tau_{Triplet}$ | Triplet relaxation |
| $K_{1211}$ | $\sigma_{Triplet}(\lambda) * \rho(\lambda)$ | Triplet absorption |
| $K_{1112}$ | $1/\tau_{Triplet-Excited}$ | Triplet excited relaxation |
| $K_{812}$ | $\Phi_{ISC}/\tau_{Triplet-Excited}$ | Reverse intersystem crossing to the single state |
| $K_{132}$ | $\Phi_{bleach-TH}/\tau_{Off}$ | Bleaching from the trans neutral |
| $K_{1311}$ | $\Phi_{Bleach-triplet}/\tau_{Triplet}$ | Bleaching from the triplet |
| $K_{1312}$ | $\Phi_{Bleach-triplet-excited}/\tau_{Triplet-Excited}$ | Bleaching from the triplet excited |

**Supplementary Table 2. List of parameters used in the simulations.**

| Parameter | Value | Reference |
| --- | --- | --- |
| $\varepsilon^{TH}_{405} = \varepsilon^{CH}_{405} = \varepsilon^{OFF}_{405}$ | 22000 $M^{-1}$ $cm^{-1}$ | [2] |
| $\varepsilon^{TH}_{488} = \varepsilon^{CH}_{488} = \varepsilon^{OFF}_{488}$ | 60 $M^{-1}$ $cm^{-1}$ | [2] |
| $\varepsilon^{T-}_{405} = \varepsilon^{C-}_{405} = \varepsilon^{ON}_{405}$ | 5260 $M^{-1}$ $cm^{-1}$ | [2] |
| $\varepsilon^{T-}_{488} = \varepsilon^{C-}_{488} = \varepsilon^{ON}_{488}$ | 61560 $M^{-1}$ $cm^{-1}$ | [2] |
| $\varepsilon^{INT}_{405}$ | 16555 $M^{-1}$ $cm^{-1}$ | Data |
| $\varepsilon^{INT}_{488}$ | 28000 $M^{-1}$ $cm^{-1}$ | Data and [2] |
| $\tau_{TH} = \tau_{CH} = \tau_{OFF}$ | 20 ps | [3] |
| $\tau_{T-} = \tau_{C-} = \tau_{INT} = \tau_{ON}$ | 1.6 ns | [4] |
| $\tau_{Rearrangement}$ | 5.1 µs | [3,5] |
| $\tau_{Deprotonation\text{-}CH}$ | 825 µs | [3] |
| $\tau_{protonation\text{-}T-}$ | 48 µs | [6] |
| $\Phi_{fluo}$ | 35 % | [2] |
| $\Phi_{TH\rightarrow CH} = \Phi_{CH\rightarrow TH}$ | 33 % | [2] |
| $\Phi_{T-\rightarrow C-} = \Phi_{C-\rightarrow T-}$ | 1.7 % | [2] |
| $\Phi_{INT\rightarrow TH}$ | 12.6 % | Data |
| $pK_a$ | 5.9 | [2] |
| $\tau_{Triplet}$ | 5 ms | [7] |
| $\tau_{Triplet\text{-}Excited}$ | 1.0 ps | [8] |
| $\Phi_{ISC}$ | 0.25 % | [7] |
| $\Phi_{RISC}$ | 0.25 % | Data |
| $\varepsilon^{Triplet}_{405}$ | 2000 $M^{-1}$ $cm^{-1}$ | [7] |
| $\varepsilon^{Triplet}_{488}$ | 10000 $M^{-1}$ $cm^{-1}$ | [7] |
| $\varepsilon^{Triplet}_{592}$ | 7500 $M^{-1}$ $cm^{-1}$ | [7] |
| $\Phi_{Bleach\text{-}TH}$ | ~ $10^{-5}$ – $10^{-6}$ % | Data |
| $\Phi_{Bleach\text{-}triplet}$ | ~ $10^{-3}$ % | Data |
| $\Phi_{Bleach\text{-}triplet\text{-}excited}$ | ~ $10^{-6}$ – $10^{-7}$ % | Data |
| $\varepsilon \rightarrow \sigma$ (conversion factor) | 3.825 x $10^{-21}$ M $cm^{-1}$ | [1] |

**Supplementary Note 2: Origin of the initial drop in rsEGFP2 photo-switching fatigue curve**

The photo-cycle of rsEGFP2 has been exhaustively studied using different spectroscopic techniques[3,5,6,9] as well as crystallographic methods[2,10]. In particular, the off-to-on transition has been characterized as a multi-step process involving an excited state isomerization reaction followed by the deprotonation of the chromophore[3,5,9]. Such a final deprotonation process results from a cascade of intermediates with different lifetimes ranging from 5 µs to 2000 µs[3,5]. Given the reported *Int* lifetimes, at moderate 405 nm energy doses (typically for RESOLFT microscopy ~ 10 – 500 W/cm$^2$), the deprotonation of the chromophore becomes the rate-limiting step of the on-switching process. Moreover, since the isomerization process is rather efficient ($\Phi_{TH \rightarrow CH}$ = 0.33[2]), an excess of UV illumination will lead to a light-induced quasi-equilibrium between the different protonated isomers of the chromophore.

We reason that the presence of such a long-lived *Int* and its large absorption cross-section at 405 nm is behind the large initial drop in the recorded fluorescence signal on the photo-switching fatigue experiments. To illustrate such an effect, we simulated the expected fluorescence response of rsEGFP2 to consecutive photo-switching cycles, just as it occurs in a fatigue experiment, and monitored the drop in the fluorescence signal from the 1$^{st}$ to the 2$^{nd}$ cycle for different deprotonation times in a 5-state model (electronic states 1-10 in Supplementary Figure 1, a simplified scheme is also included in Supplementary Figure 2a).

As the 405 nm on-switching dose is applied, the stable off-state *TH* is rapidly converted into *CH* and further on to *Int* from where it can reach the on-state, $C^-$ via a slow ground-state deprotonation. Nonetheless, the intermediate state can also undergo photo-isomerization back to *TH* by the same 405 nm dose reducing the final on-state concentration.

In general, if both on and off-states have an absorption cross-section for both on/off-switching wavelengths the cycling between the two states will never be 100%[11]. The presence of an intermediate state in the *off-to-on* transition adds another pathway that can further reduce the efficiency of the on-switching process since this state can also be excited by on/off-switching wavelengths. As shown in Supplementary Figure 2, the initial drop is non-zero for any deprotonation time and becomes more relevant around τ ~ 1 ms at 405 nm power density = 100 W/cm$^2$ and 1 ms of illumination time. By examining the time traces of the relative populations of the ground state species involved in photo-switching, we observe that a longer *Int* lifetimes reduces the concentration of molecules in the on-state at the second photo-switching cycle (orange curves in Supplementary Figure 2b), thus, yielding a lower fluorescence signal. Additionally, for a deprotonation time of 1 ms the *Int* species will accumulate at the onset of 405 nm illumination (purple curve) forming a quasi-equilibrium between *Int*, *CH* and *TH* which equilibrates around 20% of the population in the off-state, *TH* (green curve).

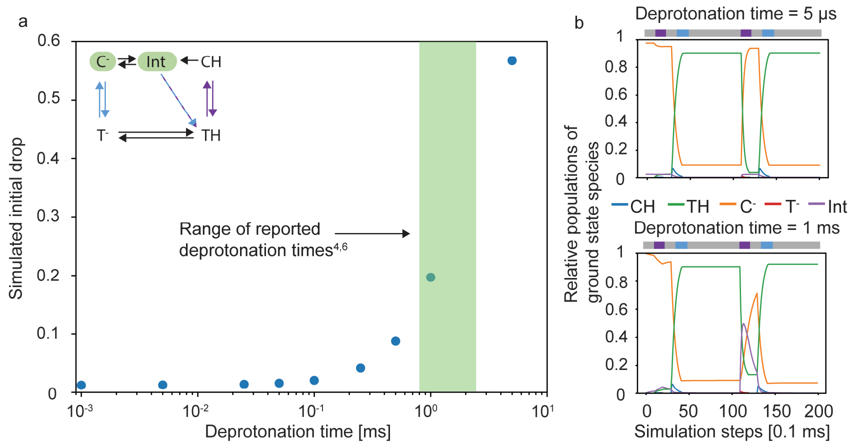


**Supplementary Figure 2. (a) Simulated drop of fluorescence between the 1$^{st}$ and 2$^{nd}$ cycles in a fatigue experiment as a function of the deprotonation time – which is analogous to the *Int* lifetime. The inset shows a schematic of the proposed photo-switching model for rsEGFP2 where $C^-$ is the fluorescent state or on-state, and *TH* is the main off-state. As the *Int* lifetime increases, the probability of a UV-light-triggered isomerization reaction from the *Int* to the off-state also increases, reducing the fluorescence observed in the second cycle. The green-shaded area shows the range of the reported deprotonation times in the literature[3,5]. The illumination doses set as input in the simulations are comparable to those of an experimental setting: 405 nm exposure for 1 ms at 100 W/cm$^2$ and 488 nm exposure for 1.2 ms at 200 W/cm$^2$. (b) Time traces of the relative populations of the ground state species involved in the photo-switching of rsEGFP2. For short deprotonation times, there is no accumulation of the *Int* population, and the majority of the *TH* concentration can be on-switched to $C^-$. On the other hand, longer deprotonation lifetimes, result in a residual fraction of the protein's ensemble population being trapped in *TH* after on-switching.**

### Supplementary Note 3: Parametrization of the intermediate species of rsEGFP2

To model the photo-switching cycle of rsEGFP2 most reliably, some of the parameters concerning the *Int* species had to be elucidated, in particular, the absorption properties at 405 and 488 nm ($\varepsilon^{INT}_{405}$ and $\varepsilon^{INT}_{488}$, respectively) as well as the $\Phi_{INT \rightarrow TH}$.

To parametrize the extinction coefficient at 488 nm, the on-switching process of rsEGFP2 was characterized with µs temporal precision using the confocal microscope described in Supplementary Note 14 and the rsEGFP2 protein embedded in a polyacrylamide (PAA) gel. The fluorophore population residing in a diffraction-limited confocal volume was firstly off-switched with a long 488 nm light dose (3 ms and ~ 40 kW/cm$^2$), immediately after, a burst of 405 nm light (5 µs and ~ 30 kW/cm$^2$) was delivered in the same volume to trigger the on-switching transition. Following a variable delay (5 µs up to 51.2 ms), another 488 nm dose (0.5 ms and ~ 40 kW/cm$^2$) was delivered to the sample and the evolution of the fluorescent signal was recorded. As shown in Supplementary Figure 3a, the immediate fluorescence response gradually increases with the delay between 405 and 488 nm pulses, reaching a plateau after a few ms. Moreover, we monitored the area below each off-switching curve and plotted them against the variable delay. In the first approximation, we pinpoint the *Int* species to be responsible for the fluorescent signal recorded after a 5 µs delay, which is around 45 % of the maximum value. Given the reported extinction coefficient of the fluorescent form, $\varepsilon^{ON}_{488}$ = 61900 M$^{-1}$ cm$^{-1}$ [2], our data suggests that $\varepsilon^{INT}_{488}$ ~ 28000 M$^{-1}$ cm$^{-1}$ which is similar to the extinction coefficient for the so-called *on fully neutral* form of rsEGFP2 found in a previous crystallographic study[2]. Additionally, we observed that for the shortest delays – 5 and 10 µs – there is a build-up of the fluorescence signal in the first few microseconds of 488 nm illumination (inset of Supplementary Figure 3b) compatible with the first relaxation step in the on-switching transition[3,5] ($CH \rightarrow Int$ in our scheme of Supplementary Figure 1) and earlier reported in time-resolved fluorescence decays in the µs time range[6].

The temporal response of the observed increase in the integrated fluorescence signal in Supplementary Figure 3b was fitted to a stretched exponential function giving a raising time of $\tau$ = 1.29 ms, in agreement with the slowest deprotonation processes resolved by spectroscopic methods[3,5]. A key assumption in the modelling of the on-switching process was that *Int* and the on-state, $C^-$, had identical spectroscopic characteristics (fluorescence lifetime and fluorescence quantum yield) except for their absorption cross-section at 405 and 488 nm, and that the transition between species leads to an increase of $\varepsilon_{488}$ and a decrease of $\varepsilon_{405}$ for $C^-$ respect to *Int* as previously reported[3,5].

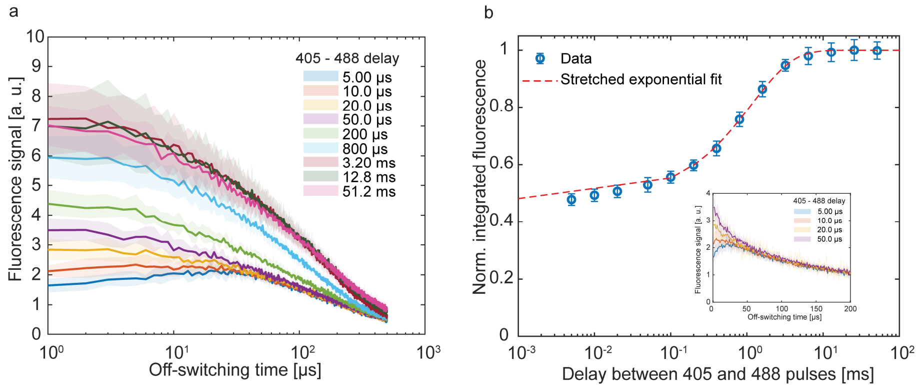


**Supplementary Figure 3. (a) Off-switching curves for multiple 405-488 delays. Longer delay between pulses correlated with an increase of the fluorescence signal peak up to a few ms when the maximum fluorescence value reached a plateau. The data shows there is a build-up of the signal in the tenths of microseconds timescale after 488 illumination for the shortest delays (5 and 10 µs). (b) Integrated fluorescence plotted against the variable delay between 405 and 488 pulses. Similarly, to (a), the area below the off-switching curve showed an exponential dependency to the delay between pulses and was fitted to a stretched exponential function with a raising time of 1.29 ms. The inset shows the building up of fluorescence in the first tenths of microseconds, suggesting the presence of a fluorescent c with such a lifetime.**

On the other hand, the $\varepsilon^{INT}_{405}$ and $\Phi_{INT \rightarrow TH}$ were estimated by fitting the photo-switching model of rsEGFP2 (electronic states 1-10 from Supplementary Figure 1) to a 405 nm on-switching ramp of increasing activation doses. The experiment consisted of consecutive photo-cycles, 405 nm to on-switch and 488 nm to off-switch and fluorescence read-out, with modulable 405 nm power densities (up to 1 kW/cm$^2$) to elucidate the fluorescence signal dependence to the on-switching dose. We decided to fix the *Int*'s lifetime during the fitting routine as such parameter has already been characterized in the literature with various methods[3,5]

As shown in Supplementary Figure 4, the observed signal increases rapidly with increasing 405 nm illumination until it reaches a plateau (around 100 W/cm$^2$), where the signal saturates. The same experiment was simulated with a 5-state photo-switching model (electronic states indexed 1-10 in Supplementary Figure 1) of rsEGFP2. No triplet state formation nor bleaching pathways were considered as each data point was an average of 9 photo-cycles and our photo-switching fatigue data does not show substantial bleaching effects after so few cycles. The output of the simulation was a modelled on-switching curve that was passed to a least-square fitting routine with $\varepsilon^{INT}_{405}$ and $\Phi_{INT\rightarrow TH}$ as the parameters to estimate. The best fit was found at $\varepsilon^{INT}_{405}$ ~ 16000 M$^{-1}$cm$^{-1}$ and $\Phi_{INT\rightarrow TH}$ ~ 12%. To our knowledge, there are no studies parametrizing the spectroscopic properties of *Int* and the values we obtained from the fit are estimations given the assumed photo-switching model. However, the evolution of the UV-vis spectra of rsEGFP2 has been characterised and from time-resolved transient absorption spectroscopy data[3], it points to a gradual decrease in the band around 405 nm indicating that the absorption in the UV region lowers as the chromophore changes from the off-state ($\varepsilon^{OFF}_{405}$ = 22000 M$^{-1}$cm$^{-1}$ [2]) to the on-state ($\varepsilon^{ON}_{405}$ = 5260 M$^{-1}$cm$^{-1}$ [2]). This behaviour is consistent with the value obtained in the fit.

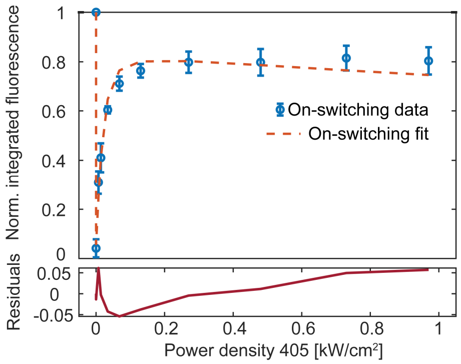


**Supplementary Figure 4. On-switching curve data and fit. Each data point is an average of 9 complete photo-cycle repetitions. The simulation consisted of a 5-state model without triplet and bleaching pathways.**

**Supplementary Table 3. Estimated parameters for the *Int* state. The value of the parameters is accompanied by the confidence interval calculated from their respective fitting routines.**

| Parameter | | Method | Estimated value | Confidence Interval (95%) |
|---|---|---|---|---|
| $\varepsilon^{INT}_{488}$ | | Time-resolved fluorescence measurement | 28000 M$^{-1}$ cm$^{-1}$ | - |
| $y = 1\text{-}\mathbf{a}\text{*}exp(\text{-}\ \mathbf{k}\text{*}t)^{\mathbf{b}}$ | ***a*** | Stretched exponential fit | 0.522 | 0.519 – 0.531 |
| | ***k*** | | 0.775 ms$^{-1}$ | 0.721 – 0.828 ms$^{-1}$ |
| | ***b*** | | 0.879 | 0.827 – 0.931 |
| $\Phi_{INT\rightarrow TH}$ | | On-switching curve fit with 5-state photo-switching model | 12.6 % | 11.7 – 13.7 % |
| $\varepsilon^{INT}_{405}$ | | On-switching curve fit with 5-state photo-switching model | 16555 M$^{-1}$ cm$^{-1}$ | 15913 – 17226 M$^{-1}$ cm$^{-1}$ |

The estimated parameters reported in Supplementary Table 3 were obtained from fitting the response of our 5-state photo-switching model to the experimental data. Nonetheless, given that such parameters are correlated and that there is an error associated to the applied 405 nm power density, other combinations of $\Phi_{INT\rightarrow TH}$ and $\varepsilon^{INT}_{405}$ would also reproduce the experimental curve with similar fidelity. We used the simulation tool to investigate such correlations among parameters by monitoring the initial drop of fluorescence between first and second photo-switching cycle.

As illustrated in Supplementary Note 2, a long-lived intermediate that can absorb 405 nm light will reduce the expected fluorescence signal at the second photo-switching cycle. The magnitude of this drop will be dictated by the 405 nm excitation dose, its spectroscopic properties ($\varepsilon^{INT}_{405}$, $\Phi_{INT\rightarrow TH}$) and the solution's pH. Given such multiparametric dependencies, there exists multiple combinations of parameters that can reproduce the same initial drop. Similarly to the model of Supplementary Figure 2, all the surface plots included in Supplementary Figure 5 correspond to a 5-state kinetic scheme without triplet state nor active bleaching pathways as they do not have a significant influence when assessing the magnitude of the initial drop, which is defined as the normalized fluorescence signal at the second cycle in a fatigue recording. The off-switching time in all the plots of Supplementary Figure 4 was 1.2 ms and the 488 nm illumination power density was 0.2 kW/cm$^2$ and $\tau_{INT}$ = 825 µs.

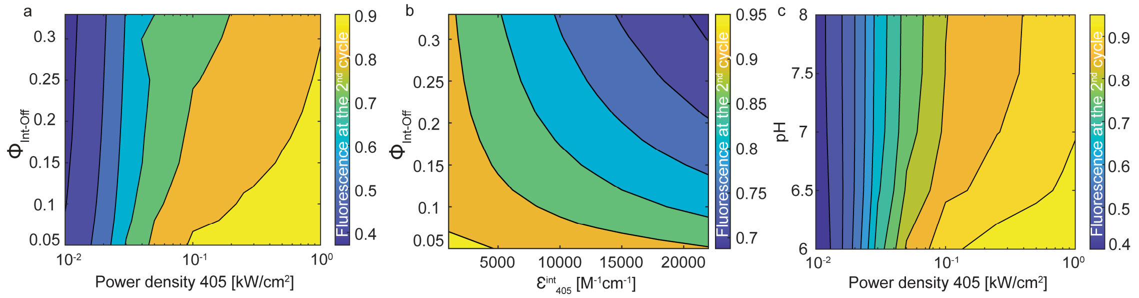


**Supplementary Figure 5. (a) Normalized fluorescence at the 2$^{nd}$ cycle as a function of the $\Phi_{INT\rightarrow TH}$ and the 405 nm irradiation power density. (b) Normalized fluorescence at the 2$^{nd}$ cycle as a function of $\Phi_{INT\rightarrow TH}$ and $\varepsilon^{INT}_{405}$. (c) Normalized fluorescence at the 2$^{nd}$ cycle as a function of the pH and the 405 nm irradiation power density.**

In the case of Supplementary Figure 5a, we examined how changing the 405 nm excitation dose affected the initial drop for a series of $\Phi_{INT\rightarrow TH}$ while keeping $\varepsilon^{INT}_{405}$ = 16000 M$^{-1}$cm$^{-1}$. From the simulation result, we observe that at low 405 nm power densities (< 50 W/cm$^2$), the illumination dose will mostly dictate the loss of fluorescence between the 1$^{st}$ and 2$^{nd}$ cycles. As we increase the dose, such loss becomes smaller as a larger share of the ensemble population is on-switched. However, an *Int* with a more efficient $\Phi_{INT\rightarrow TH}$ (~ 0.30) is expected to have a loss of fluorescence at the second cycle of around 20%, while in a less efficient one, $\Phi_{INT\rightarrow TH}$ = 0.05 the loss would be around 10%. This closely relates to the discussion on the deprotonation time (in Supplementary Note 4), a higher $\Phi_{INT\rightarrow TH}$ will result in a bigger fraction of the ensemble being redirected back to the off-state where it would be trapped and not contribute to the photo-switching turnover, yielding a lower fluorescent signal. Supplementary Figure 5a also illustrates that for a given 405 nm power density the evolution of the initial drop with $\Phi_{INT\rightarrow TH}$ is rather smooth, with many different quantum yields resulting in similar losses of fluorescence at the second cycle.

The correlation between $\Phi_{INT\rightarrow TH}$ and $\varepsilon^{INT}_{405}$ is clearly visible in Supplementary Figure 5b. For a 405 nm illumination power density of 100 W/cm$^2$, an *Int* that very efficiently isomerizes back to the *trans neutral* form (top right corner, high $\Phi_{INT\rightarrow TH}$ and $\varepsilon^{INT}_{405}$) have an expected normalised fluorescence at the second cycle around 0.7, while on the other end, one can on-switch around all the ensemble population if both $\Phi_{INT\rightarrow TH}$ and $\varepsilon^{INT}_{405}$ are rather low (left bottom corner). In between, a large family of solutions have the same expected initial drop as the two parameters appear to be anti-correlated.

In Supplementary Figure 5c, we wanted to evaluate the influence on the initial drop of two of the main experimental parameters, the power density of 405 nm and the PAA gel's pH. At low 405 nm illumination doses, the fluorescence at the second cycle is mainly determined by the excitation power density. At the experimental condition, physiological pH = 7.5, the initial drop saturates around 0.9 for power densities higher than 300 W/cm$^2$. Although the initial drop does not seem to change very rapidly as a function of the illumination power density for a given pH, Supplementary Figure 5c shows that over/underestimations of the illumination dose or the buffer's pH can lead to differences in the expected fluorescence at the second photo-switching cycle as reported recently in other green rsFPs[11].

### Supplementary Note 4: Bleaching pathways in rsEGFP2

We systematically characterized the photo-switching fatigue of rsEGFP2 by investigating the influence of the illumination doses at 405 and 488 nm. We identified two main components for the loss of fluorescence upon thousands of on/off photo-switching cycles: a fast initial drop and a gradual loss of fluorescence across hundreds/thousands of cycles that will be called fatigue fraction (from $4^{th}$ to $2000^{th}$ cycle). Moreover, we observed that mainly the latter is modulated by both illumination wavelengths, thus, we focused on studying the fatigue fraction as a function of the power density of the two illuminations. Initially, we only consider the triplet state as the main bleaching pathway for rsEGFP2 – electronic states 1-12 in the kinetic scheme of Supplementary Figure 1 -, and within this model, we identify the photo-bleaching channels to stem both from the triplet and its first excited state (electronic states 11 and 12 in Supplementary Figure 1 respectively). The addition of a photo-bleaching pathway was necessary to reproduce the 488 nm power density dependence observed in the experimental fatigue curves. Triplet state formation has traditionally been acknowledged as one of the main causes of photo-bleaching in organic dyes[12] as well as fluorescent proteins[13,14] since it can act as a reaction partner to molecular oxygen.

The experimental data points are shown in Supplementary Figure 6 below as dots (same as Figure 1d in the main text), the colour code represents the fatigue fraction (loss of fluorescence from the $4^{th}$ to the $2000^{th}$ cycle) for each combination of 405 and 488 nm power densities. Higher fatigue fraction values indicate a greater loss of fluorescence. Analogous to Figure 1d in the main text, the experimental data points are overlayed to the simulated values of the fatigue fraction across different illumination power densities for both wavelengths. In this case, the simulation only accounted for bleaching from the triplet and triplet excited states and from the colour coding it is readily observable that such a model cannot account for the increased loss of fluorescence of the experimental data at high 405 nm illumination doses. If the bleaching fraction derives only from the triplet state, the main driving force for photo-bleaching will be the 488 nm energy dose as this wavelength is preferentially absorbed by $C^-$ ($\varepsilon^{on}{}_{488} > \varepsilon^{on}{}_{405}$) from which the triplet state will appear, moreover, the triplet state has also an absorption peak at 488 nm[7] that will contribute to an enhancement of photo-bleaching. In such a model, the 405 nm dose will mainly dictate the concentration of $C^-$ and, as the available fraction of $C^-$ saturates, so will the loss of fluorescence associated with that wavelength. Our experimental data differs from the behaviour of this model, i.e. an increase of the 405 nm illumination dose produces a continuous increase of the fatigue fraction.

To simulate the loss of fluorescence of rsEGFP2 upon thousands of photo-switching cycles for different combinations of 405 and 488 nm illuminations, we first identify the effective quantum yields of the different bleaching pathways active in the model for a ramp of 488 nm illumination doses at a given 405 nm power density (dark-shaded area in Supplementary Figure 6a). Using these data points, we find a combination of effective bleaching quantum yields that reproduce well the experimental fatigue curves, and then, using the same parameters, we extrapolate to the whole two-dimensional parameter space of illumination power densities. The same procedure was carried out to simulate the fatigue fraction presented in Figure 1d in the main text, in that case however, three effective bleaching quantum yields were considered as we included a bleaching pathway from the off-state. To recreate the experimental conditions, the 488 nm off-switching time for a given power density was set such as the fluorescence signal had decayed to 20% of the initial value. Supplementary Figure 6b compares the experimental fatigue fraction (in blue) to the simulations given two different models, one considering triplet bleaching (orange dashed line) and the other considering triplet bleaching as well as bleaching from the off-state (yellow dashed line). The fatigue fraction including the off-state bleaching was slightly overestimated at low 488 nm power densities, however, the general trends of the fatigue fraction two-dimensional power density space were satisfactorily reproduced. The effective bleaching quantum yields used in the simulations are tabulated in Supplementary Table 4 below.

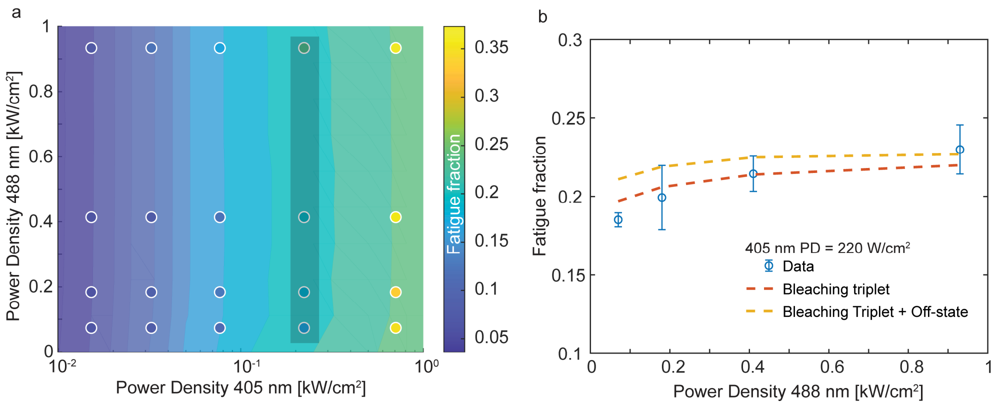


**Supplementary Figure 6. (a) Fatigue fraction data and simulation with bleaching only from the triplet state. The data (coloured dots) predicts a much higher fatigue fraction than the model at high 405 nm power densities. The data**

**points in the dark-shaded rectangle (405 nm power density = 220 W/cm$^2$) were used to fix the effective bleaching quantum yields employed in the simulations. (b) Fatigue fraction data (blue) at 220 W/cm$^2$ compared to different bleaching models: only triplet state (orange) and triplet state plus off-state (yellow).**

**Supplementary Table 4. Effective bleaching quantum yields used in the fatigue fraction simulations.**

| Bleaching model | $\Phi_{Bleach\text{-}triplet}$ | $\Phi_{Bleach\text{-}triplet\text{-}excited}$ | $\Phi_{Bleach\text{-}trans\text{-}neutral}$ |
|---|---|---|---|
| Triplet only | 1.2 x $10^{-3}$ | 0.5 x $10^{-5}$ | - |
| Triplet + Off-state | 0.8 x $10^{-3}$ | 0.2 x $10^{-5}$ | 1.0 x $10^{-5}$ |

As shown in Figure 1d in the main text, the addition of a bleaching pathway stemming from the off-state, specifically the *trans neutral* form of the chromophore, reproduced better the trends observed in the experimental data at high 405 nm power densities. We reason that the presence of the long-lived, high-absorbing *Int* increases the concentration of *TH* per cycle, and more specifically, a fraction of the protein's ensemble population (around 20 % at $\tau_{Deprotonation\text{-}C^-}$ = 1 ms, as seen in Supplementary Note 4) is trapped in *TH* during 405 nm illumination without undergoing the full on-switching transition (from *TH* to $C^-$). To assess that, we made use of the simulation tool to compute the integrated concentration of *TH** per cycle during the 405 nm illumination time as a function of the power density and the deprotonation time of the *Int* species. In this case, no bleaching channels were active as we wanted to corroborate if the presence of the *Int* species in the photo-switching contributed to an increase in 405 nm absorption from the off-state. As displayed in Supplementary Figure 7, longer deprotonation times result in up to 10-fold larger [*TH**] during 405 nm illumination per cycle at high 405 nm doses (PD > 300 W/cm$^2$). Since the isomerization reaction is quite effective, the population turnover from *TH* to *Int* happens rapidly (~ ps from *TH* to *CH* and µs from *CH* to *Int*), however from there, longer deprotonation times yield a higher probability to convert back to *TH*, hence the greater [*TH**] per illumination time each cycle. We hypothesize that such cumulative excitation of UV-light may lead to the photo-destruction of the chromophore and modelled it using an effective bleaching quantum yield from *TH**, $\Phi_{Bleach\text{-}TH}$.

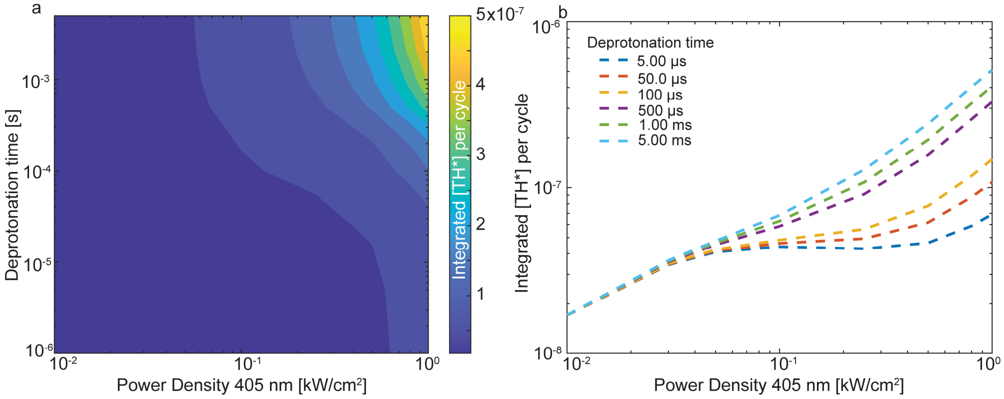


**Supplementary Figure 7. (a) Integrated concentration of *TH** per cycle during 405 nm illumination time as a function of the 405 nm power density and the deprotonation time. At the reported deprotonation time of 825 µs, the [*TH**] per cycle is nearly 10 times larger than at short deprotonation times. (b) [*TH**] per cycle as a function of the 405 nm power density for selected deprotonation times.**

We encountered variability in the observed initial drop in the photo-switching fatigue curves across different datasets. As shown in Supplementary Figure 5c, both the PAA gel's pH and the 405 nm power density will influence the magnitude of the initial drop, thus, divergences from the assumed experimental conditions may be behind the variability. Another experimental source of error might stem from the experiment procedure itself as the sample needs to be placed in focus for every measurement which implies a continuous illumination before the actual pulse scheme. Although this process is carried out at low irradiances (10 -100 µW), the variable illumination time may lead to differences in the expected response of the fluorophore. We tentatively investigated the influence of this initial perturbation on the expected initial drop for rsEGFP2 in the presence of different active bleaching channels: only from the triplet state or both the triplet and the off states. We compared the effect of initial perturbation to the expected response of a simulation without previous illumination. Note that the latter condition was the standard for all the simulations presented in the text. The perturbation was simulated by adding a pulse of both 405 and 488 nm illumination with variable length and power densities before the photo-switching fatigue curve simulation.

In Supplementary Figure 8b we simulated the expected fluorescence at the second photo-switching cycle with different bleaching pathways active in the modelling of rsEGFP2. The green dashed line in Supplementary Figure

8b represents the expected initial drop if no perturbation is applied and no bleaching channels are assumed. The input pulse scheme in the simulation was 1 ms dose of 405 nm light at 100 W/cm$^2$ and 1.5 ms of 488 nm at 150 W/cm$^2$. The green dashed line represents the maximum signal available at such illumination doses. Even if no photo-bleaching channels are active, a small perturbation of 50 mJ/cm$^2$ reduces the fluorescence output at second cycle around ~ 3% (no bleaching channels, blue curve). This reduction is enhanced further if the initial perturbation dose increases. The addition of the bleaching channels into the modelling (triplet bleaching, orange curve and triplet + off-state bleaching, yellow curve) implies a larger fluorescence drop in the 2$^{nd}$ cycle, reaching an ~ 8% fluorescence loss if all channels are active and the perturbation dose is 50 J/cm$^2$ (yellow curve) which is similar to the disagreement we observe between simulation and experimental data in Supplementary Figure 8a.

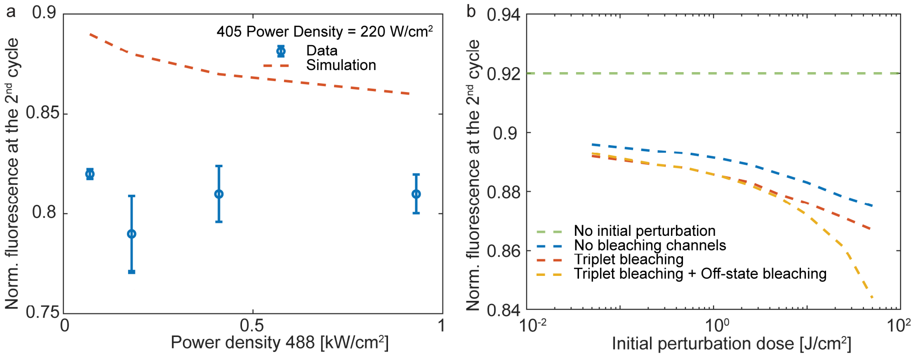


**Supplementary Figure 8. (a) Experimental data and simulated initial drop as a function of the 488 nm power density for a 405 nm power density of 220 W/cm$^2$. There is a disagreement of around 6 % between data and simulation. (b) Effect of an initial perturbation on the initial drop with different models. The simulations were carried out at 405 nm 100 W/cm$^2$ and 488 nm power density 150 W/cm$^2$ with 1.5 illumination time. If all the bleaching channels are active, the initial drop decreases by around 8 % for a 50 J/cm$^2$ energy dose.**

### Supplementary Note 5: On-switching dynamics in rsEGFP2

Complete photo-switching, i.e. moving 100% of the fluorophore's population from the on to the off state and vice versa, is in general not possible since both on and off states present non-zero absorption cross-sections for each of the wavelengths used in photo-switching[11]. In other words, the wavelength used to trigger the *off-to-on* transition can also be absorb by the on-state and promote the *on-to-off* transition equilibrating the off-state population to a small fraction after on-switching. The presence of an intermediate in the photo-cycle, like in rsEGFP2, adds a temporal dimension to this effect since the interplay between the excitation rate and the lifetime of the intermediate state will dictate how big is the fraction that equilibrates.

As introduced in Supplementary Notes 2 and 3, the presence of the long-lived intermediate state is responsible for the initial drop observed in the photo-switching fatigue experiments. This sharp decrease in the expected fluorescence of rsEGFP2 after photo-activation is a result of the light-induced quasi-equilibrium that is established between *TH, CH* and *Int* if the 405 nm illumination time is comparable to the lifetime of the intermediate state while illuminating the sample with typical 405 nm illumination intensities (~ 50 – 500 W $cm^{-2}$).

The nonlinearity of the on-switching transition also means that delivering the same total illumination dose in different manners – longer and less intense pulses rather than high-intensity bursts – yields a different on-switching efficiency. As shown in Supplementary Figure 9a below, the relative population of *TH* just before the 488 nm pulse starts decreases if the previous on-switching transition is triggered by longer pulses, while the inverse behaviour is observed in the simulation for $C^-$. While delivering the 405 nm on-switching photons slower (longer 405 nm illumination times) the probability of the *Int* transitioning back to *TH* is reduced and the turnover between the off and the on state (*TH* and $C^-$ respectively) is maximized which will result in more fluorescent photons collected during the 488 nm read-out pulse. It is important to note that the *Int* concentration before 488 nm illumination in Supplementary Figure 9a is constant and close to 0 because the simulated pulse scheme includes a long resting time (> 3 ms) between 405 and 488 nm pulses which should allow the relaxation of any *Int* population into the fluorescent state, $C^-$. On the other hand, if the on-switching dose is delivered very rapidly (< 100 µs illumination time and > 1 kW $cm^{-2}$) the concentration of *CH* after photo-activation increases. An accumulation of *CH* will lead to an increment in *TH* concentration as the isomerization reaction between these two states is very probable, shifting the equilibrium away from *Int* and, therefore, reducing the available concentration in $C^-$. When the illumination dose is delivered very slow the photo-switcher will act closer and closer to a simple on/off switch without visible effects and accumulation of intermediate states, i.e. in the right side of Supplementary Figure 9a-b the population resides almost fully in the on-state, $C^-$.

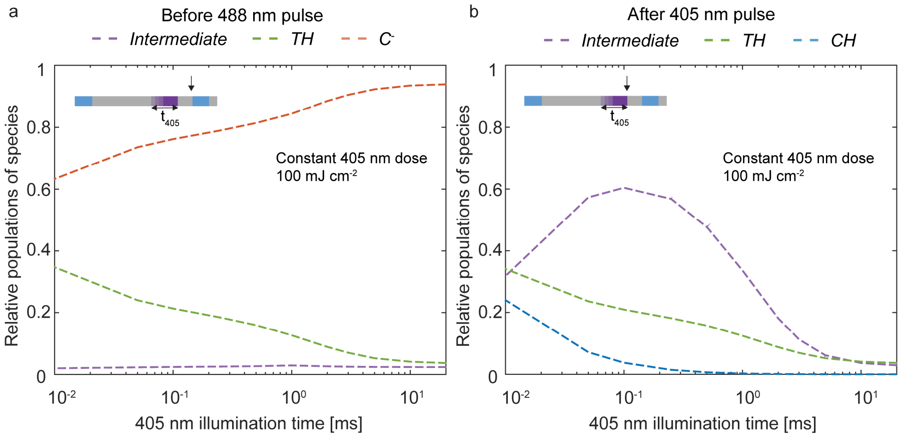


**Supplementary Figure 9. (a) Relative populations of states Int, TH and C- just before the 488 nm illumination. The simulated pulse is shown in the inset. The 405 nm illumination time was variable – from 0.01 to 20 ms – while the energy dose was kept constant at 100 mJ cm-2 – the corresponding power densities ranged from 104 to 5 W cm-2 –. The 488 nm dose was 200 W cm-2 and 1.2 ms illumination times in all the simulated points. (b) Relative populations of states Int, TH and CH just after the 405 nm illumination. The simulated pulse is shown in the inset. The 405 nm illumination time was variable – from 0.01 to 20 ms – while the energy dose was kept constant at 100 mJ cm-2 – the corresponding power densities ranged from $10^4$ to 5 W cm-2 –. The 488 nm dose was 200 W cm-2 and 1.2 ms illumination times in all the simulated points.**

**Supplementary Note 6: Parametrization of the triplet state of rsEGFP2**

To fully understand the photo-switching fatigue recovery we observed in rsEGFP2, it became apparent that we needed to include some of the triplet's spectroscopic properties. Firstly, we investigated the triplet's decaying time with µs temporal resolution in our custom-built confocal microscope using a three pulse illumination sequence, 405-488-488 nm. In these experiments, a first burst of 405 nm light (5 µs and 30 kW/cm$^2$) prepares the protein's ensemble population in the on-state, after a delay of 3 ms, a 488 nm dose (500 µs and 40 kW/cm$^2$) will off-switch the protein's population within the confocal volume. After a variable delay (from 5 µs to 500 ms) a second 488 nm dose, identical to the first one, will interrogate the same volume. The fluorescence emitted by the sample is recorded and the area below the second off-switching curve is monitored and compared to the expected fluorescence for a 0 ms delay control experiment, we call this parameter the ratiometric signal, i.e. the ratio between the integrated fluorescence signal from the second 488 nm pulse with and without a dark waiting time. As shown in Supplementary Figure 10, we observe an increase in the fluorescence signal of the second off-switching pulse as a function of the delay between the 488 nm pulses, moreover, we reason that such an increase comes from the relaxation of a dark state generated during the first 488 nm illumination pulse. Additionally, we fitted the dependence of the ratiometric signal to the delay between 488 pulses to a bi-exponential fit yielding a fast-raising time $\tau_1$ = 2.59 ms (Figure 2b in the main text) which is consistent with the reported triplet lifetime of rsEGFP2[7].

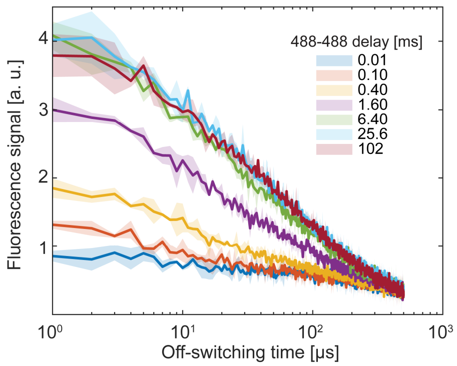


**Supplementary Figure 10. (a) Off-switching curve during the second 488 nm pulse for multiple delay times. The fluorescence signal increases with the delay until it saturates for delays longer than 6.4 ms.**

**Supplementary Table 5. Parameters of biexponential fit of Figure 2b in the main text.**

| Parameter | | Method | Estimated value | Confidence Interval (95%) |
|---|---|---|---|---|
| $y = a_1 * exp(- k_1 * t) + a_2 * exp(-k_2 * t)$ | $a_1$ | Biexponential fit | -0.995 | - 1.035 – (- 0.955) |
| | $k_1$ | | 0.386 µs$^{-1}$ | 0.358 – 0.414 µs$^{-1}$ |
| | $a_2$ | | 2.065 | 2.032 – 2.098 |
| | $k_2$ | | 7.132x10$^{-5}$ µs$^{-1}$ | -1.566x10$^{-4}$ – (1.398x10$^{-5}$) µs$^{-1}$ |

We characterised $\Phi_{RISC}$ from the triplet excited state to the singlet state from our photo-switching fatigue recovery data using a 592 nm illumination power density ramp at 488 nm illumination dose 1.5 ms and 200 W/cm$^2$. Additionally, we fixed the effective bleaching quantum yields from the all the active channels in order to reproduce the data without 592 nm illumination: $\Phi_{Bleach\text{-}triplet}$, $\Phi_{Bleach\text{-}triplet\text{-}excited}$, $\Phi_{Bleach\text{-}TH}$ which resulted in $\Phi_{RISC}$ = **0.25 %**. Comparatively, we used such reverse intersystem crossing quantum yield in another 592 nm power density ramp with higher 488 nm illumination power density (420 W/cm$^2$ and 0.9 ms illumination time) and tried to reproduce the photo-switching fatigue curves with its corresponding set of effective bleaching quantum yields. The results of the simulations are shown in Supplementary Figure 11, where the dotted lines represent the simulated photo-switching fatigue curves while the shaded areas are $\pm\sigma$ of the experimental data. Despite the experimental and modelling challenges in reproducing photo-bleaching data we believe there is a good agreement between model predictions and experiments. Overall, the output of the model is able to grasp the coarse behaviours of the data. For both experiments, the simulation tends to display a lower photo-switching fatigue recovery at lower 592 nm power densities, especially at 610 W/cm$^2$, possibly due to an underestimation of the red-shifted illumination power, nonetheless, once the effective bleaching quantum yields are fixed the simulation is consistent in reproducing the control curve and the trends suggested by the data.

It is important to note that experimental bleaching depends upon several factors regarding sample preparation, e.g. the diffusion properties and amount of oxygen dissolved in the sample. For this reason we optimized the bleaching quantum yield for each dataset in order to reproduced the observed trends in the data. The quantum yield of RISC is a molecular parameter and we argue that should be conserved in the modelling of all the experimental data.

The $\Phi_{RISC}$ value that best reproduced the trends observed in the 592 nm power density ramp is around 2-fold higher than the one reported in a study published on rsEGFP2 at 100K[7]. As mentioned, the power was measured at the objective's back aperture to calculate the power density and the laser power delivery over time was assumed constant, nonetheless, given the variability of the experimental data we estimate an error within 20 % of the measured power density. In that regard, one could explain our overestimation of the $\Phi_{RISC}$ magnitude due to an underestimation of the 592 nm power density at the sample plane.

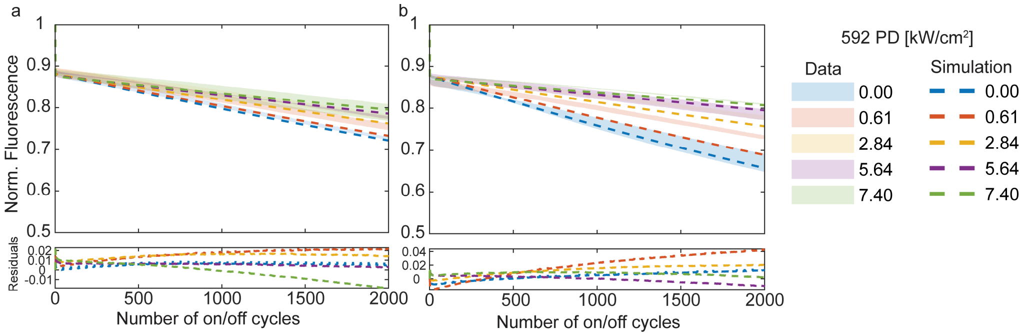


**Supplementary Figure 11. (a) Experimental data and simulation of a power ramp of 592 nm at 488 nm = 1.5 ms and 200 W/cm$^2$. The dotted lines represent the simulation while the shaded areas are ± σ of the experimental data. Below, are the residuals of each experimental curve as *data - simulation.* (b) Experimental data and simulation of a power ramp of 592 nm at 488 nm = 0.9 ms and 420 W/cm$^2$. The dotted lines represent the simulation while the shaded areas are ± σ of the experimental data. Below, are the residuals of each experimental curve as *data - simulation*.**

A similar procedure was carried out when reproducing the photo-switching fatigue data as a function of the delay between the 488 and 592 nm pulses. The parameter was set to $\Phi_{RISC}$ = **0.25 %** and a set of effective bleaching quantum yields was found to properly reproduce the data. Similarly, Supplementary Figure 12 shows a comparison of the simulations (dotted curves) and ± σ of the experimental data (shaded areas). The colour coding represents the different conditions. All the relevant parameters used in the simulations of Supplementary Figures 13 and 14 are shown in Supplementary Table 6.

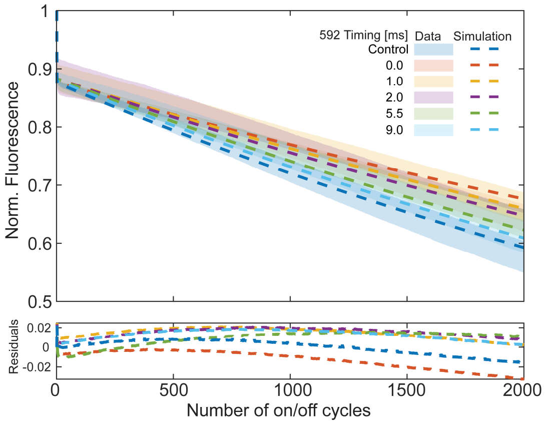


**Supplementary Figure 12. Experimental data and simulation of a power ramp of photo-switching fatigue curves with the addition of 592 nm illumination. The red-shifted pulse was incorporated into the pulse scheme with an added delay to the 488 nm dose. The dotted lines represent the simulation while the shaded areas are ± σ of the experimental data. Below, are the residuals of each experimental curve as *data - simulation.***

**Supplementary Table 6. Parameters used in the simulations of Supplementary Figures 13 and 14. Note that the illumination doses are the same as in the experiments.**

| Simulation | 405 nm dose | 488 nm dose | 592 nm dose | $\Phi_{Bleach\text{-}triplet}$ | $\Phi_{Bleach\text{-}triplet\text{-}excited}$ | $\Phi_{Bleach\text{-}TH}$ | $\Phi_{RISC}$ |
|---|---|---|---|---|---|---|---|
| **592 nm power ramp, low 488 power** | 1.0 ms | 1.5 ms | 1.0 ms | $0.8 \times 10^{-6}$ | $0.8 \times 10^{-3}$ | $1.0 \times 10^{-7}$ | 0.25 % |
| | 360 W/cm$^2$ | 200 W/cm$^2$ | 0 – 7.4 kW/cm$^2$ | | | | |
| **592 nm power ramp, high 488 power** | 1.0 ms | 0.9 ms | 1.0 ms | $1.0 \times 10^{-6}$ | $1.2 \times 10^{-3}$ | $1.0 \times 10^{-7}$ | 0.25 % |
| | 300 W/cm$^2$ | 420 W/cm$^2$ | 0 – 7.4 kW/cm$^2$ | | | | |
| **488 – 592 nm delays** | 1.0 ms | 0.9 ms | 1.0 ms | $1.0 \times 10^{-5}$ | $1.2 \times 10^{-3}$ | $5.0 \times 10^{-7}$ | 0.25 % |
| | 240 W/cm$^2$ | 300 W/cm$^2$ | 4.0 kW/cm$^2$ | | | | |

**Supplementary Note 7: Photo-switching fatigue recovery upon NIR illumination**

The light-induced recovery was measured with co-illumination in the near-infrared (NIR) spectral region by taking advantage of the Ti:Sapphire tunable laser and the confocal microscope depicted in Supplementary Figure 22. Based on a recently published work, we tuned the red-shifted co-illumination to 900 nm where the most prominent absorption peak of the triplet state of rsEGFP2 should be[7], and 810 nm, where the absorption of the triplet state for rsEGFP2 is similar to that at 592 nm. The experiments were carried out in a confocal microscope in a PAA-embedded rsEGFP2 sample and the protein was switched on and off by consecutive 405 and 488 nm pulses. As shown in Supplementary Figure 14b, the normalized light-induced recovery of the fluorescent signal after ~ 2000 photo-switching cycles was evaluated as a function of the red-shifted co-illumination power density. We observed that for power densities > 20 kW/cm$^2$ the light-induced recovery from 900 nm is significantly greater than at 810 nm which is in line with the reported triplet spectra[7]. This result suggests that the addition of 900 nm can be an efficient mechanism for photo-switching fatigue reduction, similar to the reported findings in EGFP[13]. Nonetheless, the ~ 20% light-induced recovery at 900 nm co-illumination is difficult to compare to the recovery observed at 592 nm in the widefield microscope since the spatial distribution of light-induced kinetic processes is radically different.

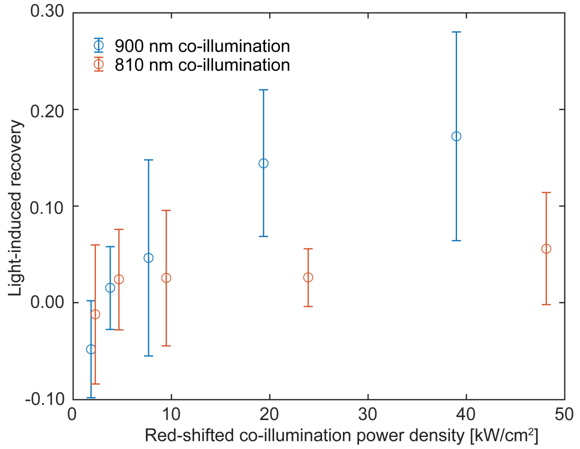


**Supplementary Figure 13. Light-induced recovery to NIR co-illumination. The effect of NIR co-illumination of photo-switching fatigue was studied in terms of the NIR power density (0 - 40 kW/cm$^2$ at 900 nm and 0 – 50 kW/cm$^2$ at 810 nm). The light-induced recovery was computed by comparing the normalized intensity after ~ 2000 photo-switching cycles with or without NIR co-illumination. For each condition, N ≥ 3 fatigue curves were taken. From these curves a mean curve per condition was calculated (with its associated standard deviation, σ). The data points correspond to the difference between the mean curve at given power density and the mean control curve after ~ 2000 photo-switching cycles. The errorbars correspond to the combined error associated with both measurements. We observe ~ 3-fold greater light-induced recovery at 900 nm than at 810 nm for similar power densities. The 405 nm dose set to 300 W/cm$^2$ and 1 ms while the laser was driven at 80 MHz with ~ 100 ps pulse width. The 488 nm dose set to 14 W/cm$^2$ and 1 ms while the laser was driven at 80 MHz with ~ 100 ps pulse width.**

**Supplementary Note 8: Photo-switching fatigue recovery upon red-shifted illumination**

The photo-switching fatigue recovery observed upon illumination with a red-shifted wavelength appears as the 592 nm dose can be efficiently absorbed by the triplet state, but not by the singlet state in addition to the non-zero probability to undergo reverse intersystem crossing. In that line, the magnitude of the photo-switching fatigue recovery induced by the red-shifted co-illumination will depend on two factors: i) how big is the photo-switching fatigue for a set of 405 and 488 illumination conditions and ii) how much of the bleaching fraction can be recovered by the 592 nm dose. For such purposes we used the simulation tool and studied the expected photo-switching fatigue recovery as a function of the illumination doses from the three wavelengths: 405, 488 and 592 nm. In the simulations, the 405 nm illumination time was kept constant at 1 ms, while the power density was fixed at 100 W/cm$^2$ when studying the dependencies of the recovery to 488 and 592 nm power densities. On the other hand, the 488 nm illumination time was tuned in order to off-switch the fluorescence by 80% when studying the dependencies of the fluorescence excitation wavelength and 592 nm, and the 488 nm illumination dose was kept at 1.2 ms and 200 W/cm$^2$ when investigating the effect of the 405 nm illumination. The bleaching parameters for the three active channels were identical to the 488-592 nm delay time simulation from Supplementary Figure 12.

The expected photo-switching fatigue recovery is relatively low at small 405 nm doses since there is no apparent bleaching, moreover, it is also noticeable that a threshold of ~ 2 kW/cm$^2$ of 592 nm power density should be surpassed to have approximately 5% of fatigue recovery. The red-shifted wavelength co-illumination recovery is favored at higher 405 nm power densities (> 200 W/cm$^2$) when the $C^-$ *concentration* is maximum, and high 592 nm energy doses as well, since more absorption per illumination dose will occur. The magnitude of the recovery, however, evolves slower as we move towards the upper right corner of the plot in Figure 2i in the main text (high 405 and 592 nm power densities) as the off-state bleaching becomes more dominant and the triplet concentration saturates.

Similarly, we also investigated the magnitude of the fatigue recovery as a function of the 488 and 592 nm power densities as displayed in Figure 2j in the main text. The photo-switching fatigue recovery is below 5 % until 592 nm power density ~ 1 kW/cm$^2$ for low 488 nm doses. The recovery is enhanced at a high 488 nm power density, although it reaches a plateau > 600 W/cm$^2$ for 592 nm power densities > 10 kW/cm$^2$ which may indicate the excitation saturation of the triplet state with red-shifted co-illumination. As the energy dose (illumination time * illumination power density) is roughly constant throughout the range of power densities investigated, the adverse effects from the triplet's high absorption at 488 nm are diminished and the magnitude of the recovery is mainly driven by the triplet state 592 nm excitation probability.

The importance of the light-induced photo-switching fatigue recovery is readily observable if one attains to the results shown in Figure 1g in the main text where the fatigue diminishes as the total pulse length increases. In that experiment, the illumination doses for all the different studied pulses were the same and only the dark time (no illumination of the sample) was successively increased. By increasing the pulse length of an order magnitude, the photo-switching fatigue was reduced by ~ 18 % after 1600 on/off cycles, however, if such a pulse scheme were implemented in an imaging context the temporal resolution would be heavily compromised.

Supplementary Figure 14a compares two approaches for fatigue recovery after 1600 cycles: increasing pulse length and co-illumination with 592 nm light. The horizontal lines in Supplementary Figure 14a represent the normalized fluorescence intensity recovery by increasing the waiting time per pulse compared to the standard 10 ms pulse, while the blue dots show the recovery experience by co-illuminating with 592 nm in a standard 10 ms dwell time. Notably, co-illumination with a red-shifted wavelength achieves similar fatigue recovery to longer pulses but preserves temporal resolution. These results demonstrate that while longer pulses effectively reduce photo-switching fatigue, co-illumination provides a more practical solution for imaging by minimizing bleaching effects without sacrificing time resolution.

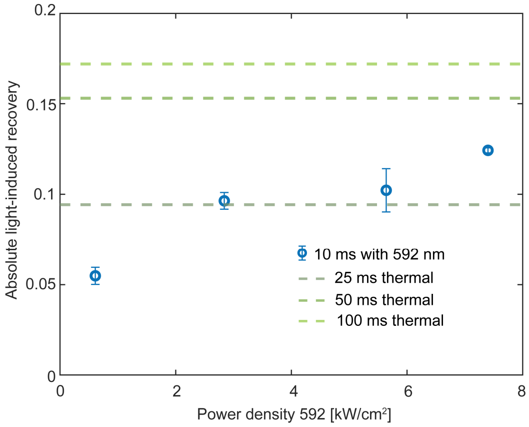


**Supplementary Figure 14. Comparison of light-induced fatigue recovery to thermal fatigue recovery from Figure 1g. The horizontal lines correspond to the difference of the normalized intensity after 1600 cycles for a given pulse length to the standard 10 ms pulse. Longer pulse lengths yielded greater recovery. In blue, is the experimental data of fatigue recovery after 1600 cycles for a 488 nm power density of 420 W/cm$^2$.**

**Supplementary Note 9: Photo-switching fatigue of other green negative photo-switchers**

To test the generalizability of this mechanism in other green negative photo-switchers, we investigated the fatigue of different rsFPs as well as their response to the 592 nm co-illumination. We characterized 4 other negative photo-switchers: rsEGFP(N205S)[15], a mutant closely related to rsEGFP2[16]; Dronpa(M159T), a green RSFP from Anthozoan origin[17]; and two variants of the rsGreen family[18], rsGreen1 and rsGreenF.

As shown in Figure 3d in the main text, the maximum recovery is observed for the rsGreen family proteins, especially notable for rsGreenF where it is > 10 % for a 592 nm power density ~ 4 kW/cm$^2$. In Supplementary Figure 15, we display the raw photo-switching fatigue curves from which the plot in Figure 3d was built. Each RSFP presents a different fatigue fingerprint, for instance, rsEGFP(N205S) showed a very prominent fatigue with no recovery at all. We hypothesize that the differences we observed between the photo-switching fatigue fingerprints of the two proteins may originate from i) different triplet bleaching channels balance and ii) the smaller off-switching quantum yield of N205S leads to a higher triplet absorption of 488 nm per illumination cycle – a similar parallelism can be drawn between rsGreen1 and rsGreenF, being the former the slower photo-switcher –. Concerning i), it is shown in Figure 2h in the main text how the balance in the relative strength of the bleaching channels from the triplet and triplet excited states greatly modulates the recovery response of rsEGFP2. If triplet excited bleaching is the dominant contribution, the photo-switching fatigue is greater given that the triplet state has a prominent absorption peak around 490 nm[7], moreover, no recovery is expected when that becomes the dominant bleaching pathway. Given that the states involved in the photo-cycle might be conserved among rsEGFP(N205S) and rsEGFP2, a skewed balance towards the triplet excited bleaching pathway in the former appears as a plausible explanation for the differences observed in their red co-illumination behaviour.

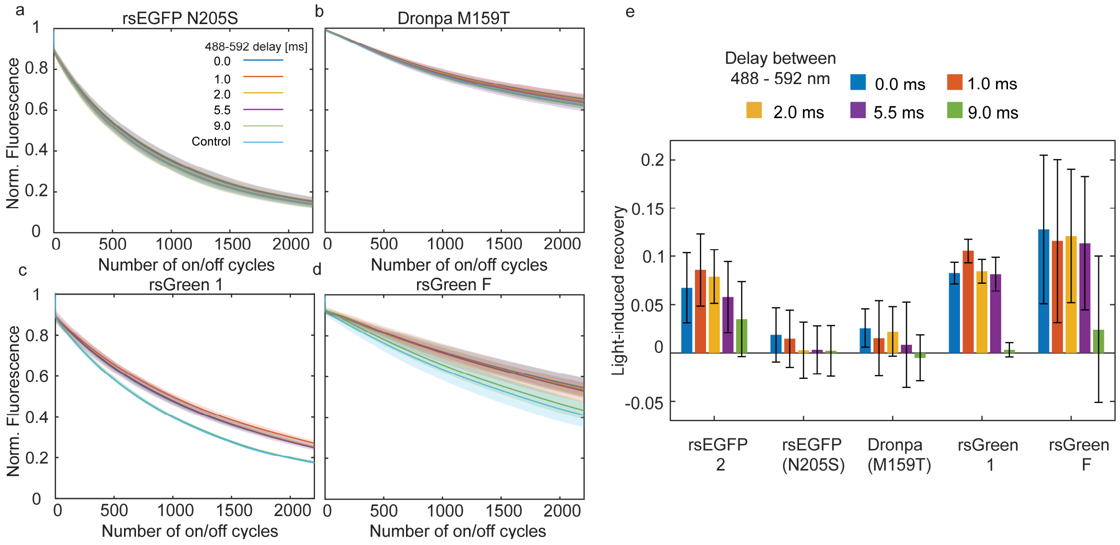


**Supplementary Figure 15. (a) rsEGFP(N205S) photo-switching fatigue curve as a function of the 488-592 nm delay time. (b) Dronpa M159T photo-switching fatigue curve as a function of the 488-592 nm delay time. (c) rsGreen 1 photo-switching fatigue curve as a function of the 488-592 nm delay time. (d) rsGreen F photo-switching fatigue curve as a function of the 488-592 nm delay time. (e) Summary of the light-induced recovery, i.e. the normalized fluorescence signal recovered after 2000 on/off cycles, for the 5 RSFPs included in the study. The colour code corresponds to the delay between 488 and 592 nm pulses. For each condition, N ≥ 3 fatigue curves were taken. From these curves a mean curve per condition was calculated (with its associated standard deviation, σ). The data points correspond to the difference between the mean curve at given power density and the mean control curve after 2000 photo-switching cycles. The errorbars correspond to the combined error associated with both measurements. The shaded areas in (a)-(d) represent ± σ associated with each mean curve for every condition.**

Additionally, we investigated how the photo-switching fatigue responded to adding a dark waiting time after the 488 nm illumination. Note that the minimum time camera read-out time for a 30x30 µm$^2$ FOV was 1.5 ms, therefore, that was set as the shortest dark time. As shown in Supplementary Figure 16, the response of the different proteins followed the trends displayed in the previous figure. Noticeably, rsEGFP(N205S) did not show any improvement in the photo-switching fatigue upon an increase in the dwell time of the photo-switching cycle, unlike the rest of the rsFPs screened in this study. To explain this, we reason that the small off-switching quantum yield of rsEGFP(N205S) linked to the high absorption of the triplet state leads to the rapid photo-bleaching displayed in the protein. The fact that rsEGFP(N205S) is such a slow photo-switcher contributes to the photo-bleaching since a higher energy dose – 488 nm power density x illumination time – is necessary to bring a significant amount of the ensemble population to the off-state (80-90 %), moreover, the relative probability to populate the triplet state rather than off-switching is higher than in a faster photo-switcher like rsEGFP2, thus, making the 488 nm absorption by the triplet state a more probable. This also could explain why rsGreen1 – a slower variant of the rsGreen family compared to rsGreenF – exhibits a substantially higher photo-switching fatigue. In summary, we argue that in slower photo-switchers, the light delivered to off-switch has a higher chance of bleaching from triplet states (ground and excited) than in other faster switchers.

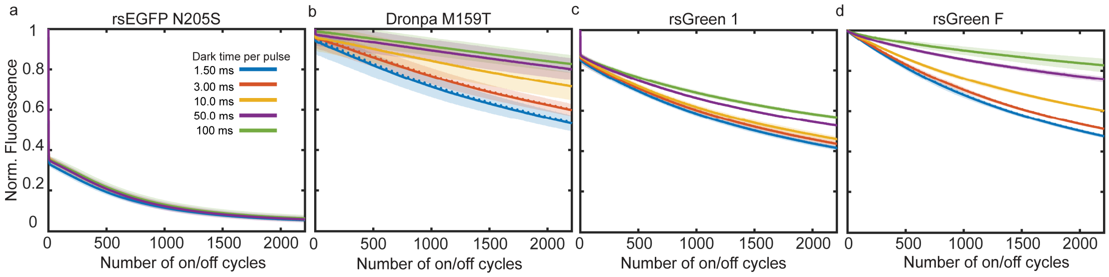


**Supplementary Figure 16. (a) rsEGFP(N205S) photo-switching fatigue curve as a function of the dark waiting time. (b) Dronpa M159T photo-switching fatigue curve as a function of the dark waiting time. (c) rsGreen 1 photo-switching fatigue curve as a function of the dark waiting time. (d) rsGreen F photo-switching fatigue curve as a function of the dark waiting time. For each condition, N ≥ 3 fatigue curves were taken. From these curves a mean curve per condition was calculated (with its associated standard deviation, σ). The shaded areas represent ± σ associated with each mean curve for every condition.**

**Supporting Note 10: Imaging multiplexing with RSFPs at high-spatiotemporal resolution**

In recent years, RSFPs have been incorporated into imaging multiplexing strategies that rely on their differentiated light-driven kinetics to unmix and recover the identity of specifically labelled structures in biological imaging. Approaches such as TMI[19] or LIGHTNING[20] carefully resolve the off-switching in time to assess the identity of the fluorophore, while strategies like exNEEMO[21,22] locate the different proteins in a multidimensional unmixing space based on their expected emission given different levels of 405 nm photo-activation. It is important to note that the better separation between different RSFPs in all these methods occurs at low excitation power densities, when the characteristic off-switching time between species are more distinct. This becomes crucial in approaches which rely on a camera-based detection system – like TMI[19] and exNEEMO[21,22] – that imposes a low time-resolution limit ~ 1 ms.

To examine how good the kinetic-based unmixing is at higher excitation power densities, we simulated the behaviour of 4 rsEGFP2-like RSFPs with different off-switching quantum yields (from $0.1*QY_{rsEGFP2}$ to $1*QY_{rsEGFP2}$) as a function of the 488 nm intensity. For each simulated fluorophore, the off-switching curve was simulated (with 2% gaussian noise added to mimic experimental conditions) and fitted with a monoexponential decay function to extract the characteristic time, $\tau_{fluorophore}$. The $\tau_{fluorophore}$ of each simulated fluorophore was subtracted to the extracted $\tau_{rsEGFP2}$ at the same 488 nm power density to obtain the $\Delta\tau$ between each simulated RSFP and rsEGFP2. The results of the simulations are shown in Supplementary Figure 17, below. At lower power densities ( < 100 $W/cm^2$) the separation between species is larger than the time-resolution of the camera making it possible to distinguish them from rsEGFP2, however, as we approach the typical 488 nm power densities for high-spatiotemporal resolution imaging (green-shaded area, 0.1 – 2 $kW/cm^2$) the difference between the characteristic times shrinks and we are rapidly below the camera time-resolution limit cut-off (red-shaded area, < 1 ms). Intuitively, one can see that a method such as TMI[19] that depends on identifying the proteins based on their off-switching fingerprint is not possible to implement at these 488 nm power densities if the target RSFPs to unmix have similar off-switching kinetics.

Other methods such as exNEEMO[21,22] while not relying on resolving the off-switching process can see a decrease in the accuracy of the unmixing since the fluorescence emission of negative photo-switchers is linked to the off-switching kinetics. If two RSFPs only differ in their off-switching quantum yields, such as in the simulation of Supplementary Figure 17, their relative positions in the unmixing space will be determined by the integral below the off-switching curve. Therefore, if the off-switching process of both RSFPs is similar enough (and all other spectroscopic properties are equal between the RSFPs) so it will be the emitted fluorescence.

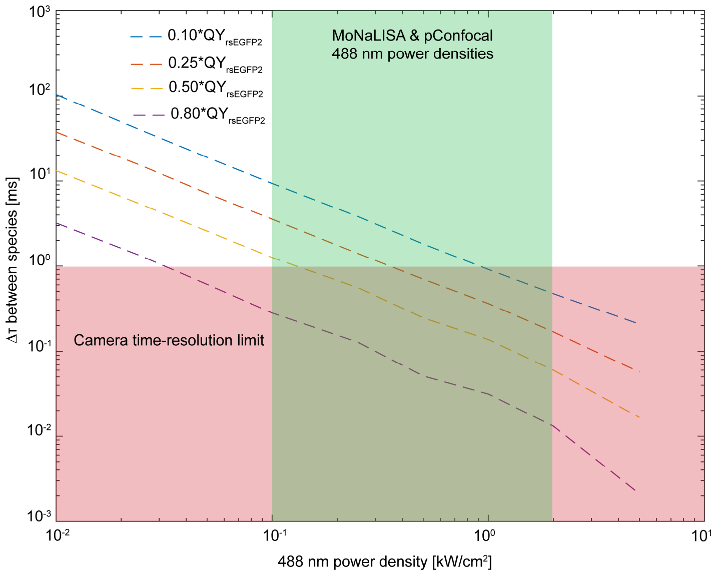


**Supplementary Figure 17. Difference between off-switching characteristic times as a function of the 488 nm power density. The off-switching of 5 different rsEGFP2-like fluorophores was simulated at multiple power densities with 2% gaussian noise. Each RSFP has a different off-switching quantum yield (from 0.1*QYrsEGFP2 to 1*QYrsEGFP2) and the characteristic off-switching time, τfluorophore, is extracted by fitting a monoexponential decay function. Each τfluorophore is compared to τrsEGFP2, to compute the difference Δτ in ms. In the green-shaded area we show the range of power densities used typically in high-spatiotemporal resolution imaging. When Δτ falls below the camera time-resolution cut-off, the proteins are not easily distinguishable only relying on the camera's time-resolution.**

**Supplementary Note 11: Photo-bleaching recovery with 592 nm co-illumination in live-cell imaging**

To quantify the effect of the red-shifted co-illumination in a live-cell imaging context, we monitored the photo-bleaching experienced in mitochondria expressing rsEGFP2 labelling the outer-membrane-protein 25 (OMP-25) imaged in parallelized confocal mode. In each image, the integrated intensity of 4 regions of interest (ROIs) was calculated – 1 of the ROIs was used to measure the image background and the remaining 3 ROIs contained the labelled structure – and monitored across the 500 frames of the time-lapse recordings. From the intensity traces, we obtained the photo-bleaching profiles for each image which were fitted with a monoexponentially decaying function, and as a result, we obtained the bleaching constant for every image. In Supplementary Figure 18a, we show the bleaching constant for every image as a function of their signal-to-background ratio in the first frame of the acquisition. We observed that adding the 592 nm co-illumination slows down the photo-bleaching experienced by the mitochondria (larger bleaching constants in Supplementary Figure 18a). In that sense, the red-shifted co-illumination prolonged the time-lapse by tenths of frames in recordings showing similar initial signal-to-background ratios proving to be a powerful method for minimizing photo-bleaching in samples where the brightness of the cellular structures of interest is limited by the transfection efficiency.

We further tested the effect of a red-shifted wavelength co-illumination in the observed photo-bleaching of live-cells in a super-resolution imaging context, specifically, RESOLFT microscopy within the MoNaLISA modality. We evaluated the photo-bleaching recovery effect by comparing the normalised fluorescence signal after 20 imaging frames with and without the secondary illumination in vimentin and actin endogenously tagged with a set of rsFPs. To build the respective bleaching curves, the signal from two sub-areas – one corresponding to the structure, one corresponding to the dark background, each of ~ 8×8 $\mu m^2$ – of the MoNaLISA FOV (~ 40×40 $\mu m^2$) was monitored across the time-lapse recording. The counts per area of each region are calculated and the background is subtracted on a frame-by-frame basis, afterwards, the signal is normalised to the first frame. The results are extracted from an average of 3-5 cells (or curves) per condition.

As we observed in the PAA gels for rsEGFP2 the recovery effect is greater when a large share of the protein's ensemble population has undergone both 405 and 488 nm excitations, therefore, we added the 592 nm illumination to the 488 nm patterned illumination path. With this optical design, we made sure that the 592 nm multi-foci pattern was co-aligned to both 405 and 488 nm multi-foci patterns. We tested the recovery effect from the red-shifted co-illumination in vimentin labelled with rsEGFP2 in HeLa cells as a function of the irradiation power density with an analysis pipeline as described above. The results displayed in Supplementary Figure 18b show a decrease in the bleached fraction after 20 imaging frames as the 592 nm illumination power density increases.

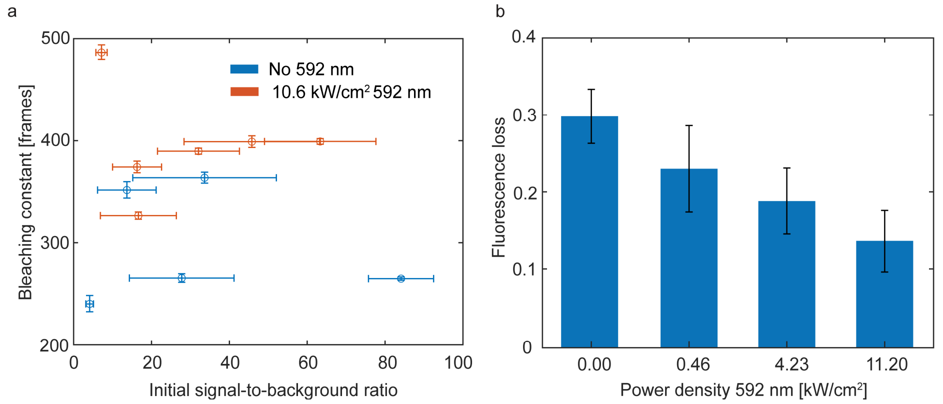


**Supplementary Figure 18. (a) Bleaching constant as function of the initial signal-to-background ratio. Larger bleaching constant indicates slower bleaching. At similar initial signal-to-background ratio, the addition of 592 nm co-illumination extends the available number of frames. (b) Bleached fraction after 20 MoNaLISA frames for different 592 nm illumination power densities in vimentin tagged with rsEGFP2 in HeLa cells. The bleached fraction is reduced as the 592 nm power density increases.**

**Supplementary Note 12: Photo-toxicity assessment in live-cell imaging context**

To assess if the 592 nm co-illumination led to any adverse effects in the imaged cells, we monitored the behaviour of mitochondria under parallelized confocal illumination conditions. In particular, we manually annotated the occurrence of events that report on the mitochondrial network mobility. Such events include branching, visualized as transient deformations of the mitochondrial membrane; fission; fusion and long stretching/retraction, which appear as transient and pronounced elongation or retraction of the mitochondrial tubules. Examples of such events are shown in Supplementary Figure 19a. The frame number when the events occurred was noted down as a timestamp for all the observed events in all the images with and without 592 nm co-illumination and summed together in bins of 25 frames. In total, we observed 224 such events in 5 images without 592 nm and 321 events in 6 images with 592 nm co-illumination. From these pool events, the cumulative probability distribution was computed as shown in Supplementary Figure 19b. Our data suggests that the addition of the 592 nm co-illumination did not affect the mobility of the mitochondrial network as represented by the counted events as both datasets, with and without 592 nm, show a similar distribution across time. Moreover, we did not see changes in the mitochondrial network morphology typically associated with photo-toxicity such as swelling or blobbing.

Additionally, we investigated whether the 592 nm co-illumination had adverse effects on the cells that were being imaged. For that purpose, we used the DNA repairing protein XRCC1 (X-Ray cross complementary factor 1)[23] as a quantitative reporter for light-induced damage in the cell's nucleus. XRCC1 acts as a central loading platform for DNA repair[23] and it manifests as a bright puncta in the nuclei. In our photo-damage assay, we counted the bright puncta in the nucleus of different cells (N = 8-10) before and after taking a MoNaLISA image with different 592 nm co-illumination power densities. As shown in Figure 20c, the additional red-shifted illumination did not incur an increase in light-induced DNA damage compared to a normal MoNaLISA imaging recording.

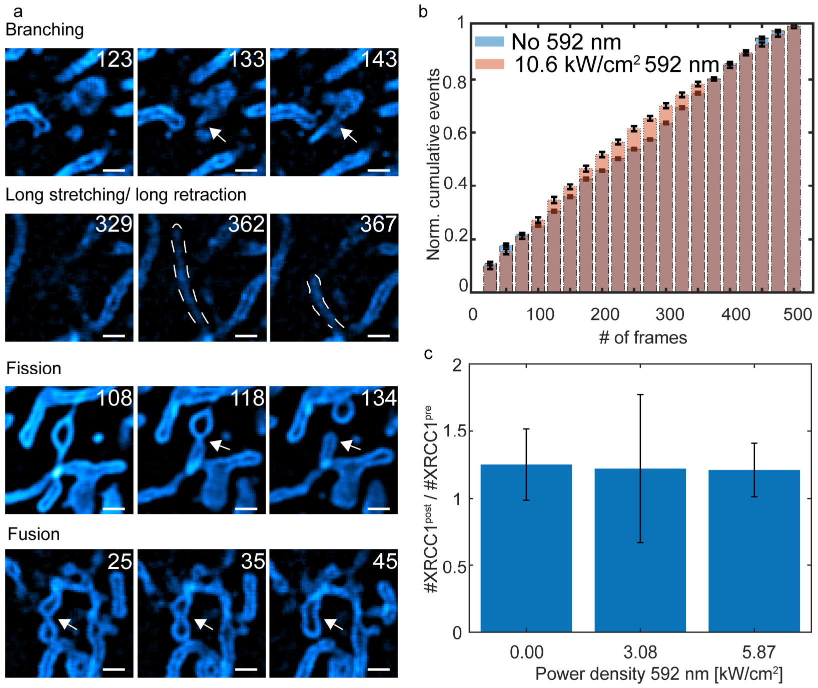


**Supplementary Figure 19. (a) Examples of reported events in the mitochondria datasets. The scale bars are 1 µm. (b) Cumulative probability distribution of the observed events across the 500 frames of the parallelized confocal recording. In total, 224 events were annotated in 5 images without 592 nm co-illumination and 321 events in 6 images with 592 nm co-illumination. (c) Light-induced DNA damage assay. The bright puncta on the cell nucleus from DNA damage reporter XRCC1 were counted before and after taking a MoNaLISA image with different 592 nm power densities. The addition of the red-shifted wavelength didn't lead to an increase in light-induced DNA damage.**

**Supplementary Note 13: Optical set-ups**

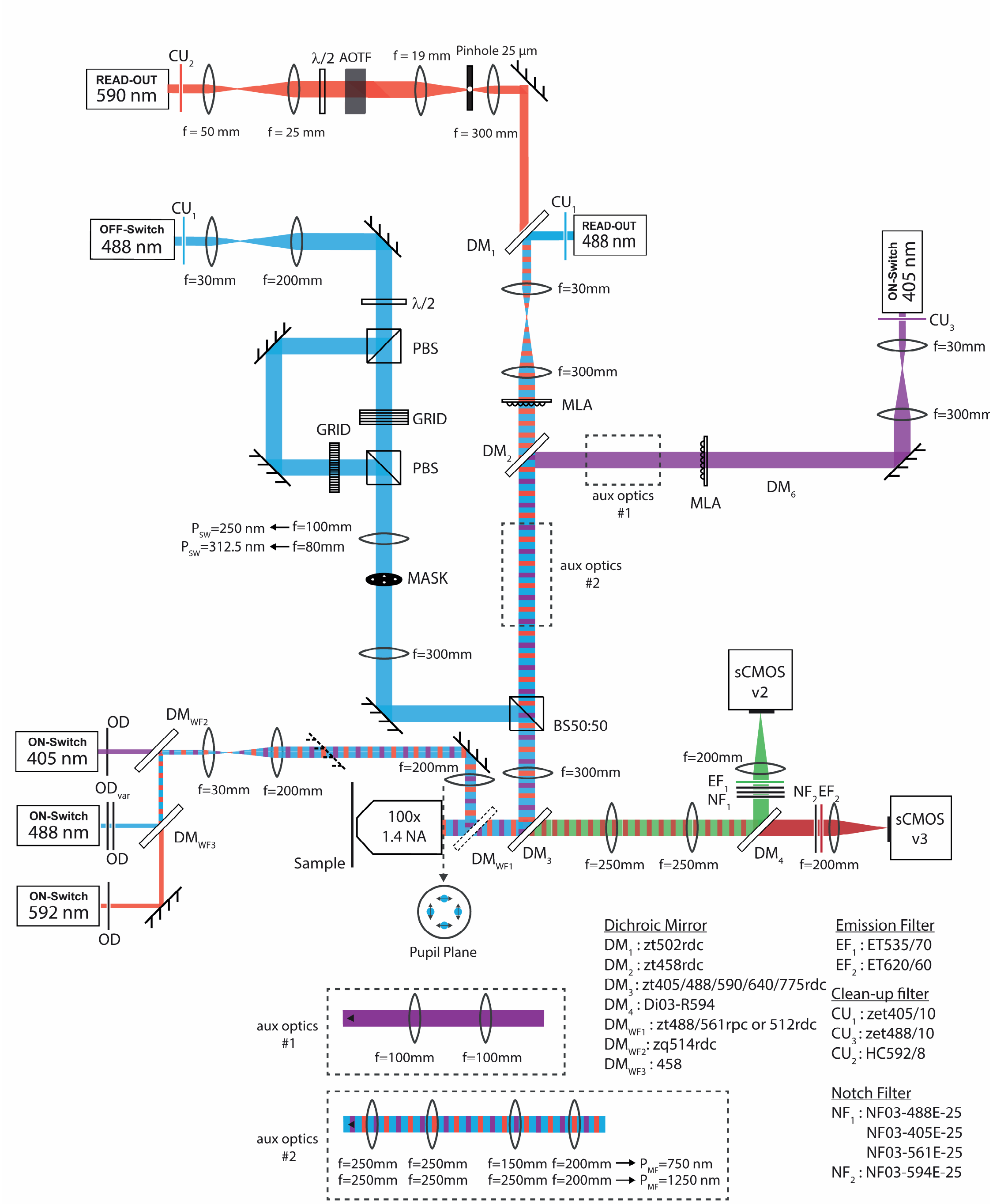


**Supplementary Figure 20. Widefield (WF) microscope set-up. The photo-switching fatigue of all proteins was carried out on this microscope as well as the RESOLFT imaging. Two pairs of microlenses array (MLA) are incorporated in the optical paths to create the multi-foci pattern at the sample plane. These optical elements were removed from the set-up when using the WF mode for photo-switching fatigue characterization. The off-switching path was only used in imaging to create the confinement necessary to achieve sub-diffractive resolution.**

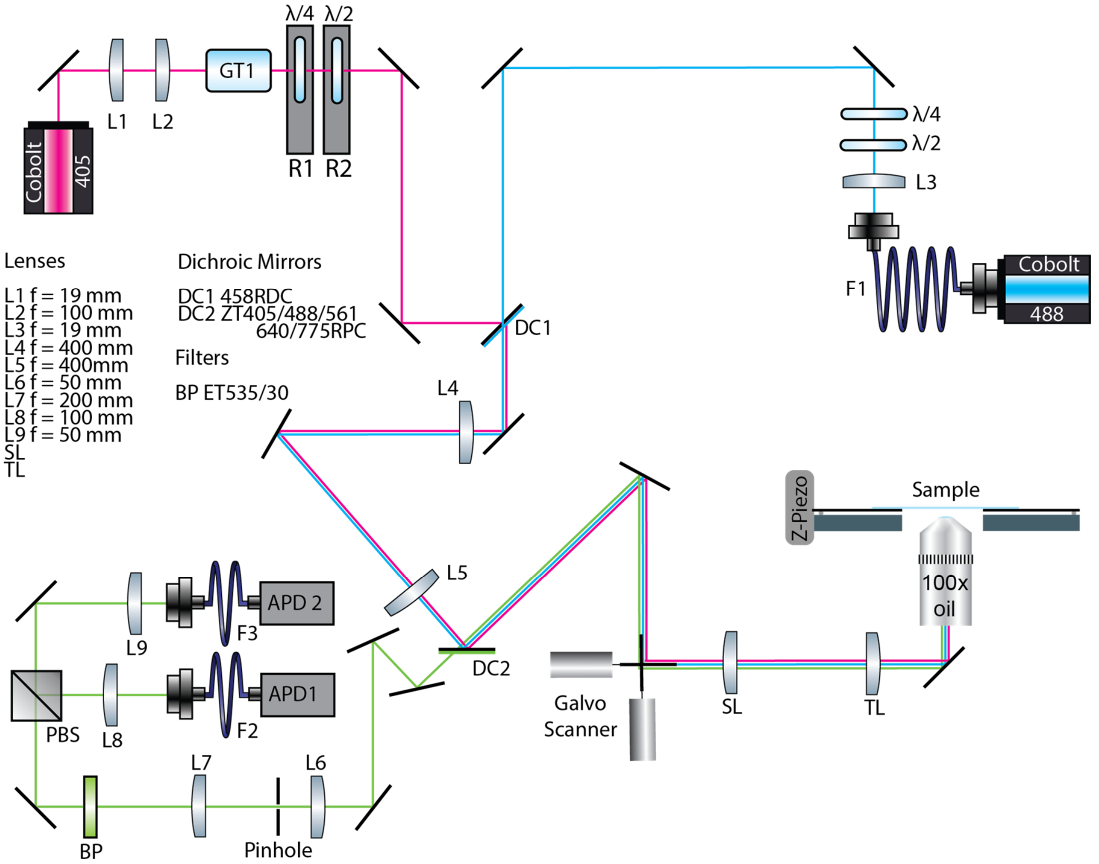


**Supplementary Figure 21. Point-scanning microscope set-up. This microscope was employed for the characterization experiments that required higher temporal resolution. The microscope incorporates two point-detectors with high temporal resolution. The signal from both detectors was summed to create the final signal. Elements not listed in the Figure are listed below. Fiber Optics: F2: FG105LGA (Thorlabs), F3: AFS50 (Thorlabs). Polarization rotators: R1 and R2: K10CR1/M (Thorlabs), Polarization Optics: GT1: GTH10M-A (Thorlabs), λ/4: 460-680 achr. (B. Halle Nachfl, Berlin, Germany), λ/2: 460-680 achr. (B. Halle Nachfl), PBS: CCM2- PBS251/M (Thorlabs). Lenses: SL: 50 mm (Leica Microsystems, Wetzlar, Germany), TL: 200 mm (Leica). Scanners: Galvo Scanner XY: 6215H Galvanometric mirrors + 71215HHJ Servo Driver (Cambridge Technology, Bedford, MA, USA), Piezo Stage Z: LT-Z-100 (Piezoconcept, Lyon, France).**

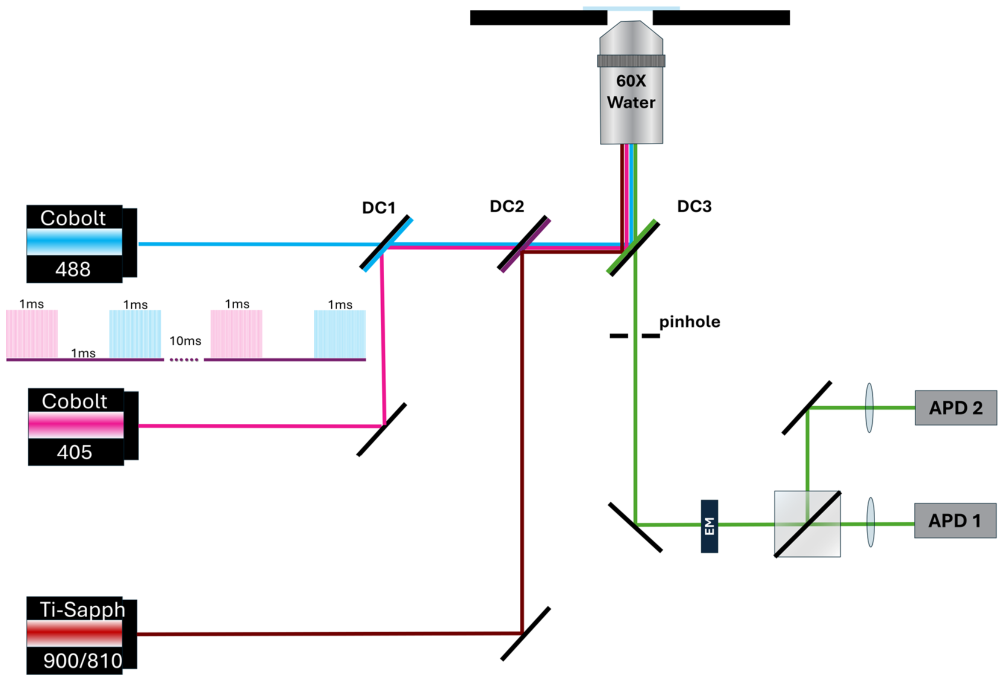


**Supplementary Figure 22. Near-infrared confocal microscope set-up. This microscope was employed for the characterisation experiments in the NIR region. The red-shifted co-illumination was tuned employing the Ti:Sapphire laser. The microscope incorporates two point-detectors with high temporal resolution. The signal from both detectors was summed to create the final signal.**

**Supplementary Note 14: Photo-physical simulation of rsEGFP2**

To characterize the behaviour observed in the photo-switching experiments we developed a kinetics simulation tool that computes the time-evolution of the emissive state concentration according to a network of interconnected electronic states. The simulation tool was developed as Python script and solves analytically the system of linear ordinary differential equations that describe the photo-physics of the fluorophore. The theoretical framework is briefly described in this note.

Being $C_i$ represent the concentration of species *i*, the general set of rate equations for a system having *j* such species will be defined as

$$\frac{dC_i}{dt} = \sum_j k_{ij} C_j \quad (1)$$

where $k_{ij}$ is the kinetic constant for the reaction that transforms the *j-th* species into the *i* species when $j \neq i$. The kinetic constant $k_{ii}$, represents the depopulation of the *i* species according to a kinetic scheme. It is calculated by summing all the kinetic rates of transitions reducing the concentration of *i*

$$k_{ii} = -\sum_{j \neq i} k_{ji} \quad (2)$$

In general, to describe the concentration evolution of state *i* within a network of *j* interconnected states one must solve a set of differential equations such as (1) for each of the involved species. In that regard, the problem can be expressed in matrix form and simplifies to

$$\frac{d\boldsymbol{C}(t)}{dt} = \boldsymbol{KC} \quad (3)$$

All the kinetic rates connecting states in the system are collected in matrix **K** in equation 3. In that sense, the constant describing the $i \rightarrow j$ reaction will be indexed in position $\boldsymbol{k_{ji}}$. Similarly, vector **C** contains the concentrations for each state with the same indexing as **K**. The solution of the differential equation in the matrix form is analogous to its scalar equation and yields,

$$\boldsymbol{C}(t) = \exp(\boldsymbol{K}t)\, \boldsymbol{C}(t = 0) \quad (4)$$

Being **C** *(t =0)* the set of initial concentrations for each species. From here, calculating the matrix exponential requires solving an eigenvalue problem[24], since

$$\exp(\boldsymbol{K}t) = \boldsymbol{U}\exp(\boldsymbol{\lambda}t)\boldsymbol{U}^{-1}, \qquad \text{with } \boldsymbol{K} = \boldsymbol{U\lambda U}^{-1} \quad (5)$$

where **λ** is the diagonal eigenvalues matrix and **U** is the eigenvectors matrix. Therefore, to obtain the concentration at any time point for a given fluorophore one should construct a suitable kinetic matrix **K** with its photo-physical parameters and consider a reasonable set of initial conditions.

Generally, the kinetic constants connecting the different species within the scheme will represent spontaneous/thermally induced processes (non-light-induced) or light-induced processes. Since the probability of the latter is modulated by the irradiation intensity, the kinetic matrix needs to include the photon fluxes for each illumination dose within the pulse scheme. To practically emulate any experiment, we represent the experimental pulse scheme as a concatenation of so-called kinetic windows. For every kinetic window, the status of all the illumination wavelengths in the pulse scheme are defined – on or off –, as well as the photon flux delivered to the sample during that time. The different lasers are simulated as square waves, and two simplified detection modes are implemented: time-resolved or integrated signal.

**Supplementary Table 7. Parameters used in the characterization experiments with the different rsFPs.**

| Experiment | 405 nm dose | 488 nm dose | Red-shifted dose |
|---|---|---|---|
| **On-Switching curve (Supplementary Figure 4)** | 1 ms | 2.0 ms | - |
| | 0 - ~ 1 kW/cm$^2$ | 120 W/cm$^2$ | - |
| **592 nm power ramp, low 488 power (Figure 2e, Supplementary Figure 11a)** | 1.0 ms | 1.5 ms | 1.0 ms |
| | 360 W/cm$^2$ | 200 W/cm$^2$ | 0 – 7.4 kW/cm$^2$ |
| **592 nm power ramp, high 488 power (Figure 2e, Supplementary Figure 11b)** | 1.0 ms | 0.9 ms | 1.0 ms |
| | 300 W/cm$^2$ | 420 W/cm$^2$ | 0 – 7.4 kW/cm$^2$ |
| **488 – 592 nm delays rsEGFP2 (Figure 2f, Supplementary Figure 12)** | 1.0 ms | 0.9 ms | 1.0 ms |
| | 240 W/cm$^2$ | 300 W/cm$^2$ | 4.0 kW/cm$^2$ |
| **Photo-switching fatigue characterization (Figure 1b-d,f, Supplementary Figure 6a,b)** | 1 ms | 2.5 – 0.5 ms | - |
| | 15 – 700 W/cm$^2$ | 70 -930 W/cm$^2$ | - |
| **Thermal recovery rsEGFP2 (Figure 1g, Supplementary Figure 14)** | 1 ms | 5 ms | - |
| | 240 W/cm$^2$ | 40 W/cm$^2$ | - |
| **Confocal 405 – 488 nm delays (Supplementary Figure 3a,b)** | 5 µs | 500 µs | - |
| | 30 kW/cm$^2$ | 40 kW/cm$^2$ | - |
| **Confocal 488 – 488 nm delays (Figure 2b, Supplementary Figure 10)** | 5 µs | 500 µs | - |
| | 30 kW/cm$^2$ | 40 kW/cm$^2$ | - |
| **On-switching doses with constant total energy rsEGFP2 (Figure 2a)** | 20 – 0.3 ms | 1 ms | - |
| | 5 – 300 W/cm$^2$ | 200 W/cm$^2$ | - |
| **488 – 592 nm delays rsEGFP(N205S) (Figure 3a,d, Supplementary Figure 15a)** | 1 ms | 3.7 ms | 1 ms |
| | 240 W/cm$^2$ | 200 W/cm$^2$ | 4.16 kW/cm$^2$ |
| **488 – 592 nm delays Dronpa(M159T) (Figure 3a,d, Supplementary Figure 15b)** | 1 ms | 0.6 ms | 1 ms |
| | 240 W/cm$^2$ | 220 W/cm$^2$ | 4.00 kW/cm$^2$ |
| **488 – 592 nm delays rsGreen1 (Figure 3a,d, Supplementary Figure 15c)** | 1 ms | 1.5 ms | 1 ms |
| | 240 W/cm$^2$ | 190 W/cm$^2$ | 3.75 kW/cm$^2$ |
| **488 – 592 nm delays rsGreenF (Figure 3a,d, Supplementary Figure 15d)** | 1 ms | 1.0 ms | 1 ms |
| | 240 W/cm$^2$ | 240 W/cm$^2$ | 4.27 kW/cm$^2$ |
| **Thermal recovery rsEGFP(N205S) (Supplementary Figure 16a)** | 1 ms | 1.5 ms | - |
| | 7 W/cm$^2$ | 2.4 kW/cm$^2$ | - |
| **Thermal recovery Dronpa(M159T) (Supplementary Figure 16b)** | 1 ms | 0.6 ms | - |
| | 80 W/cm$^2$ | 220 W/cm$^2$ | - |
| **Thermal recovery rsGreen1 (Supplementary Figure 16c)** | 1 ms | 2.0 ms | - |
| | 240 W/cm$^2$ | 110 W/cm$^2$ | - |

| | | | |
|---|---|---|---|
| **Thermal recovery rsGreenF (Supplementary Figure 16d)** | 1 ms | 1.0 ms | - |
| | 240 W/cm$^2$ | 240 W/cm$^2$ | - |
| **NIR power ramp (Figure 2g, Supplementary Figure 13)** | 80 MHz for 1 ms | 80 MHz for 1 ms | ~ 50 s |
| | 300 W/cm$^2$ | 14 kW/cm$^2$ | 0 – 50 kW/cm$^2$ |

**Supplementary Table 8. Parameters used in the imaging experiments with the different rsFPs.**

| Experiment | 405 nm dose | 488 nm dose | 592 nm dose |
|---|---|---|---|
| **Photo-switching fatigue in vimentin-rsEGFP2 in HeLa cells (Supplementary Figure 18)** | 0.5 ms | 1.7 ms Off-Switching<br>1 ms Read-out | 1.0 ms |
| | 200 W/cm$^2$ | 1.2 kW/cm$^2$ Off-Switching<br>0.5 kW/cm$^2$ Read-out | 0 – 11.2 kW/cm$^2$- |
| **Photo-switching fatigue in LifeAct-rsEGFP2 in U2OS cells (Figure 4c,d)** | 0.5 ms | 1.7 ms Off-Switching<br>1 ms Read-out | 1.0 ms |
| | 200 W/cm$^2$ | 1.2 kW/cm$^2$ Off-Switching<br>0.5 kW/cm$^2$ Read-out | 0 & 11 kW/cm$^2$ |
| **Photo-switching fatigue in LifeAct-rsEGFP(N205S) in U2OS cells (Figure 4d)** | 0.5 ms | 4 ms Off-Switching<br>1 ms Read-out | 1.0 ms |
| | 75 W/cm$^2$ | 1.12 kW/cm$^2$ Off-Switching<br>0.71 kW/cm$^2$ Read-out | 4.8 kW/cm$^2$ |
| **Photo-switching fatigue in LifeAct-Dronpa(M159T) in U2OS cells (Figure 4d)** | 0.5 ms | 1 ms Off-Switching<br>1 ms Read-out | 1.0 ms |
| | 120 W/cm$^2$ | 1.12 kW/cm$^2$ Off-Switching<br>0.71 kW/cm$^2$ Read-out | 6.5 kW/cm$^2$ |
| **Photo-switching fatigue in LifeAct-rsGreen1 in U2OS cells (Figure 4d)** | 0.5 ms | 1.3 ms Off-Switching<br>1 ms Read-out | 1.0 ms |
| | 120 W/cm$^2$ | 1.2 kW/cm$^2$ Off-Switching<br>0.71 kW/cm$^2$ Read-out | 4.8 kW/cm$^2$ |
| **Photo-switching fatigue in LifeAct-rsGreenF in U2OS cells (Figure 4d)** | 0.5 ms | 1.3 ms Off-Switching<br>1 ms Read-out | 1.0 ms |
| | 75 W/cm$^2$ | 1.12 kW/cm$^2$ Off-Switching<br>0.71 kW/cm$^2$ Read-out | 4.8 kW/cm$^2$ |
| **DNA damage assay (Supplementary Figure 19c)** | 0.5 ms | 1.7 ms Off-Switching<br>1 ms Read-out | 1.0 ms |
| | 200 kW/cm$^2$ | 1.2 kW/cm$^2$ Off-Switching<br>0.5 kW/cm$^2$ Read-out | 0 - ~ 6.0 kW/cm$^2$- |
| **Photo-switching fatigue in OMP-25-rsEGFP2 in U2OS cells (Figure 4a, Supplementary Figure 18a, Supplementary Figure 19a,b)** | 0.5 ms | 1 ms | 1.0 ms |
| | 100 W/cm$^2$ | 0.5 kW/cm$^2$ Read-out | 0 & 10.6 kW/cm$^2$ |
| **Multiplexing of 4 RSFPs in parallelized confocal (Figure 3b,c)** | 0.5 ms / 3 ms | 5 ms | - |
| | 100 W/cm$^2$ | 0.9 kW/cm$^2$ | - |
| **Unmixing with WF + Imaging in parallelized confocal (Figure 4b)** | 0.5 ms | 5 ms | 1.0 ms |
| | 350 W/cm$^2$ | 0.9 kW/cm$^2$ | 10.6 kW/cm$^2$ |

### REFERENCES


1. Tkachenko, N. V. *Optical Spectroscopy: Methods and Instrumentations*. (Elsevier Science, 2006).
2. El Khatib, M., Martins, A., Bourgeois, D., Colletier, J.-P. & Adam, V. Rational design of ultrastable and reversibly photoswitchable fluorescent proteins for super-resolution imaging of the bacterial periplasm. *Sci. Rep.* **6**, 18459 (2016).
3. Woodhouse, J. *et al.* Photoswitching mechanism of a fluorescent protein revealed by time-resolved crystallography and transient absorption spectroscopy. *Nat. Commun.* **11**, 741 (2020).
4. Testa, I., D'Este, E., Urban, N. T., Balzarotti, F. & Hell, S. W. Dual Channel RESOLFT Nanoscopy by Using Fluorescent State Kinetics. *Nano Lett.* **15**, 103–106 (2015).
5. Uriarte, L. M. *et al.* Structural Information about the *trans* -to- *cis* Isomerization Mechanism of the Photoswitchable Fluorescent Protein rsEGFP2 Revealed by Multiscale Infrared Transient Absorption. *J. Phys. Chem. Lett.* **13**, 1194–1202 (2022).
6. Volpato, A. *et al.* Extending fluorescence anisotropy to large complexes using reversibly switchable proteins. *Nat. Biotechnol.* **41**, 552–559 (2023).
7. Rane, L. *et al.* Light-Induced Forward and Reverse Intersystem Crossing in Green Fluorescent Proteins at Cryogenic Temperatures. *J. Phys. Chem. B* **127**, 5046–5054 (2023).
8. Klán, P. & Wirz, J. Photochemistry of Organic Compounds: From Concepts to Practice. in (2009).
9. Coquelle, N. *et al.* Chromophore twisting in the excited state of a photoswitchable fluorescent protein captured by time-resolved serial femtosecond crystallography. *Nat. Chem.* **10**, 31–37 (2018).
10. Grotjohann, T. *et al.* rsEGFP2 enables fast RESOLFT nanoscopy of living cells. *eLife* **1**, e00248 (2012).
11. Bourges, A. C. *et al.* Quantitative determination of the full switching cycle of photochromic fluorescent proteins. *Chem. Commun.* **59**, 8810–8813 (2023).
12. Ringemann, C. *et al.* Enhancing Fluorescence Brightness: Effect of Reverse Intersystem Crossing Studied by Fluorescence Fluctuation Spectroscopy. *ChemPhysChem* **9**, 612–624 (2008).
13. Ludvikova, L. *et al.* Near-infrared co-illumination of fluorescent proteins reduces photobleaching and phototoxicity. *Nat. Biotechnol.* (2023) doi:10.1038/s41587-023-01893-7.
14. Byrdin, M., Duan, C., Bourgeois, D. & Brettel, K. A Long-Lived Triplet State Is the Entrance Gateway to Oxidative Photochemistry in Green Fluorescent Proteins. *J. Am. Chem. Soc.* **140**, 2897–2905 (2018).
15. Chmyrov, A. *et al.* Nanoscopy with more than 100,000 'doughnuts'. *Nat. Methods* **10**, 737–740 (2013).
16. Grotjohann, T. *et al.* Diffraction-unlimited all-optical imaging and writing with a photochromic GFP. *Nature* **478**, 204–208 (2011).
17. Ando, R., Flors, C., Mizuno, H., Hofkens, J. & Miyawaki, A. Highlighted Generation of Fluorescence Signals Using Simultaneous Two-Color Irradiation on Dronpa Mutants. *Biophys. J.* **92**, L97–L99 (2007).
18. Duwé, S. *et al.* Expression-Enhanced Fluorescent Proteins Based on Enhanced Green Fluorescent Protein for Super-resolution Microscopy. *ACS Nano* **9**, 9528–9541 (2015).
19. Qian, Y., Celiker, O. T., Wang, Z., Guner-Ataman, B. & Boyden, E. S. Temporally multiplexed imaging of dynamic signaling networks in living cells. *Cell* **186**, 5656-5672.e21 (2023).
20. Chouket, R. *et al.* Extra kinetic dimensions for label discrimination. *Nat. Commun.* **13**, 1482 (2022).
21. Valenta, H. *et al.* Separation of spectrally overlapping fluorophores using intra-exposure excitation modulation. *Biophys. Rep.* **1**, 100026 (2021).
22. Valenta, H. *et al.* Per-pixel unmixing of spectrally overlapping fluorophores using intra-exposure excitation modulation. *Talanta* **269**, 125397 (2024).
23. Serebrovskaya, E. O. *et al.* Light-induced blockage of cell division with a chromatin-targeted phototoxic fluorescent protein. *Biochem. J.* **435**, 65–71 (2011).
24. Berberan-Santos, M. N. & Martinho, J. M. G. The integration of kinetic rate equations by matrix methods. *J. Chem. Educ.* **67**, 375 (1990).
